# Self-powered piezo-bioelectronic device mediates tendon repair through modulation of mechanosensitive ion channels

**DOI:** 10.1101/2020.08.03.227868

**Authors:** Marc A. Fernandez-Yague, Alex Trotier, Sunny Akogwu Abbah, Aitor Larrañaga, Arun Thirumaran, Aimee Stapleton, Syed A. M. Tofail, Matteo Palma, Abhay Pandit, Manus J Biggs

**Affiliations:** National University of Ireland Galway, CÚRAM, Centre for Research in Medical Devices, H91W2TY, Galway, Ireland; University of the Basque Country, Department of Mining-Metallurgy Engineering and Materials Science & POLYMAT, Barrio Sarriena, 48013 Bilbao, Spain; University of Limerick, Department of Physics, V94 T9PX, Ireland; Queen Mary University of London, Materials Research Institute and School of Biological and Chemical Sciences, Mile End Road, London E1 4NS, UK

## Abstract

Tendon disease constitutes an unmet clinical need and remains a critical challenge in the field of orthopaedic surgery. Innovative solutions are required to overcome the limitations of current tendon grafting approaches, and bioelectronic therapies are showing promise in the treatment of musculoskeletal disease, accelerating functional recovery through the activation of tissue regeneration signalling pathways (guided regeneration). Self-powered bioelectronic devices, and in particular piezoelectric materials represent a paradigm shift in biomedicine, negating the need for battery or external powering and complementing existing mechanotherapy to accelerate the repair processes. Here, we show the dynamic response of tendon cells to a piezoelectric collagen-analogue scaffold comprised of aligned nanoscale fibres made of the ferroelectric material poly(vinylidenefluoride-co-trifluoroethylene), (PVDF-TrFE). We demonstrate that electromechanical stimulation of tendon tissue results in guided regeneration by ion channel modulation. Finally, we show the potential of the bioelectronic device in regulating the progression of tendinopathy associated processes using a rat Achilles tendinopathy model. This study indicates that body motion-powered electromechanical stimulation can control the expression of TRPA1 and PIEZO2 receptors and stimulate tendon-specific tissue repair processes.

## Introduction

Tendon disorders, including tendinopathy, trauma-induced injury or age-related degeneration, are commonly the result of athletic or repetitive activity^1–3^. In the US alone, more than 102.5 million adults are affected by tendon disorders every year^4^. Tendon injuries typically occur in the Achilles and the rotator cuff tendons, commonly with partial or total rupture occurring within the mid-substance or enthesis^5^. Following injury, disorganised tissue function and deposition can lead to the formation of scar tissue, proteoglycan accumulation and calcification, resulting in poor biomechanical properties and impaired function. Critically, tendon disorder mismanagement places a considerable burden on healthcare systems (> $2 billion annually and post-surgery complications result in nearly 1 million additional days of inpatient care each year). Thus, effective tendon repair strategies present a significant unmet clinical need^4,6^.

Tendon is a dynamic specialised tissue that undergoes continuous remodelling and adaption to mechanical loading and unloading during physical activity. Indeed, many recent studies confirm that resident tendon cell populations are highly mechanosensitive and are responsible for orchestrating the repair processes after injury^7–10^. To accomplish this, cells rely on membrane receptor connections between the microenvironment and intracellular proteins, which initiate signalling pathways to modulate the cellular response. under physiological conditions, tendon cells are subjected to constant mechanical deformations of 3% to 4% strain that mediates tendon homeostasis^7,11^. Changes to these strain conditions modulate cellular catabolic or anabolic activities, resulting in the structural reorganisation of the tendon tissue^12–16^. For example, studies have shown that excessive tendon loading can initiate fibroblast proliferation and promote the synthesis of mediators of inflammatory cascades, leading to increased collagen deposition along with the orientation of stress^17^. Conversely, concentric or eccentric exercise or ultrasound therapy has been extensively studied and clinically used as an approach to enhancing tendon regeneration^18–20^.

As well as responding to mechanical stimulation, an often-overlooked property of tendon tissue is its intrinsic piezoelectricity (Fig. 1). Piezoelectricity or electromechanical coupling is associated with highly oriented collagen structures, that can result in polypeptidic molecules with short-range crystallinity^21,22^ (Fig. 1b). The piezoelectric properties of dried collagenous tissues, including Achilles tendon, were first reported by Fukada *et al.*^23^ Subsequently, electromechanical coupling in wet conditions was shown by Ikushima *et al.,* who demonstrated stress-induced polarisation of Achilles tendon when acoustically stimulated in the MHz range^24^. At the cellular level, bioelectrical signalling has been shown to coordinate cell activity and endogenous electrical fields are critical in mediating cell migration and in inducing successful wound healing^25–27^. The effect of mechanically-induced electrical cues on non-excitable cells and tissue repair processes, however, remains elusive.

**Fig. 1.**
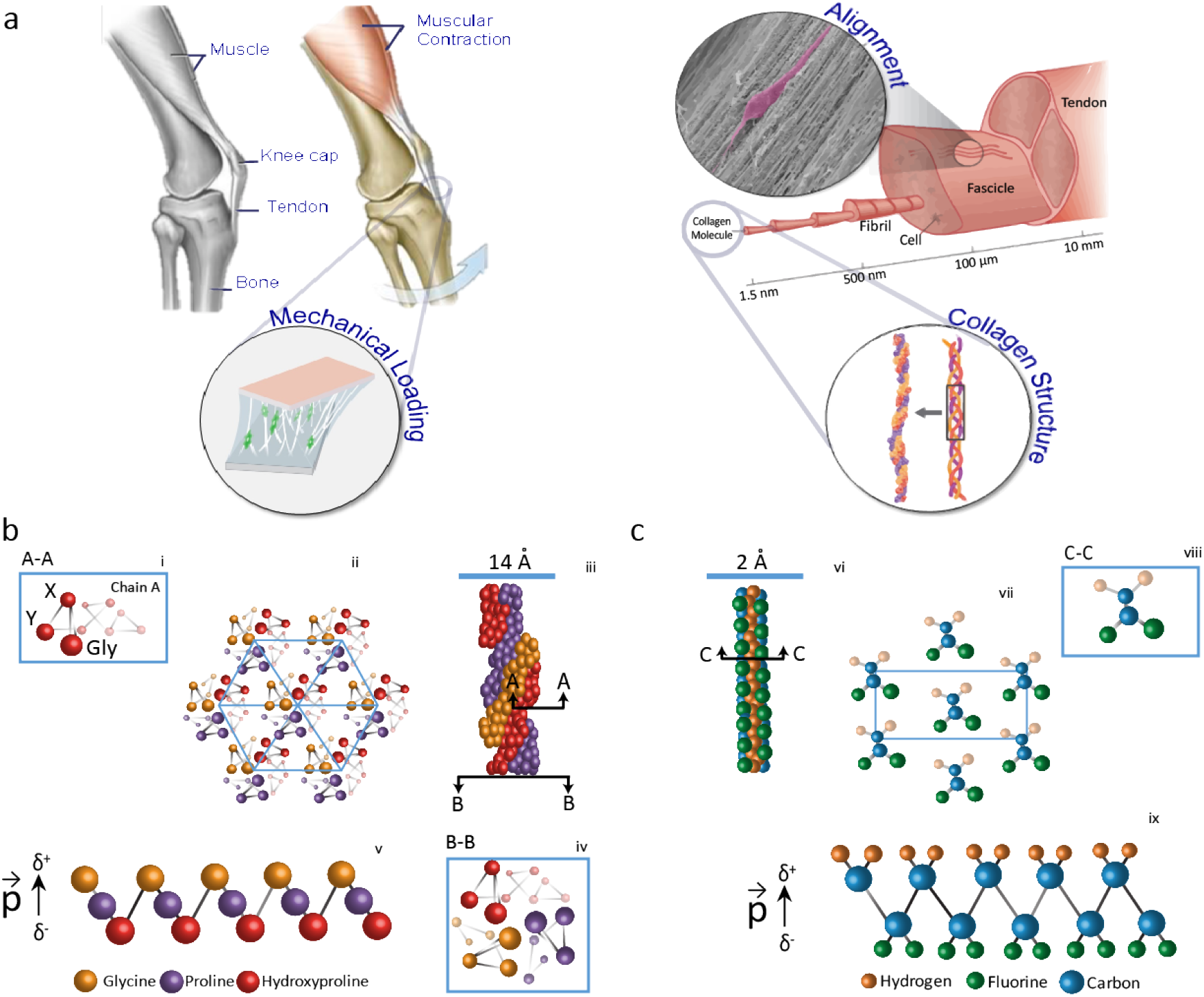
Overview of the tendon electromechanical environment and the piezoelectricity origin analogy of collagen and PVDF-TrFE fibres. Tendon is a dynamic tissue that connects muscle to bone and is under continuous mechanical loading and unloading. **a**, This mechanical stress is borne by a highly organised extracellular matrix composed of collagen type I, high-tensile strength and piezoelectric material which undergoes electrical polarization in response to mechanical loading. **b**. Schematic representation of the crystal structure of the collagen triple helix consisting of three polypeptides chains stabilised via hydrogen bonding. **i** A-A: Cross-sectional schematic of an individual alpha-chain showing the triple Gly (glycine) -X (proline) -Y (hydroxyproline) residues. Polar bonds between carbonyl (CO groups) and amine (peptide NH group of glycine residues) groups along the backbone of collagen results in a dipole moment or electrical polarisation. **iii** Crosssection of multiple individual collagen molecules resulting in a non-centrosymmetric hexagonal fibril arrangement **iv** B-B: A cross-section of the collagen triple-helix core, here the three helical chains and glycine residues are observable. **v** Representation of individual electric dipole **c**, **vi** Representation of the PVDF-TrFE all-trans (TTTT) zigzag planar configuration top view. **vii** Cross-section of the crystal and along the orthogonal axis of the all-trans (TTTT) zigzag planar configuration **viii** Representation of individual electric dipole combining of fluorine (green sphere), carbon (blue sphere), and hydrogen (orange sphere). PVDF-TrFE has a non-centrosymmetric and dipoles generated by the high electronegative difference between hydrogen and fluorine atoms **ix** Representation of the chain molecular structure of electroactive β-phase of PVDF-TrFE showing a resultant dipole moment.

Externally applied electrical stimulation (ES) is currently being used clinically as an adjunct therapy to the treatment of fracture non-unions, in the promotion of spinal fusion, for pain relief and to promote collagen synthesis in skin and ligament/tendon tissues ^28–31^. Pulsed electromagnetic field (PEMF) and capacitive coupling (CC) approaches to ES are non-invasive but generate a weak electrical current (<1 μA), the units are often heavy and large making patient compliance a concern and the dermal electrodes can produce skin irritations and infections. Additionally, clinical evidence that supports the clinical efficacy of PENF or CC has been inconclusive. Conversely, direct current (DC) ES approaches have shown clinical efficacy and offer higher levels of stimulating electrical currents (10-100 μA), shown to promote stem cell differentiation during bone healing. Furthermore, in the last 30 years, many studies have conclusively demonstrated the clinical efficacy of DC stimulators (Zimmer Biomet SpF and OsteoGen) (BGS) in improving the success rate of spinal fusion procedures and non-healing fractures^32,33^. Critically however, a significant roadblock to the clinical translation of ES therapies has arisen from a lack of effective, safe and non-invasive technologies that do not require the use of penetrating microneedles, components which have been shown to raise the risks of infection and are associated with unspecific stimulation of off-target tissues^4,34,35^. The need for more efficient and less invasive electrically stimulating systems is forcing a paradigm shift in the biomedical engineering field, where novel bioelectronic strategies that control somatic cell functions and enhance specific tissue regenerative process are desirable^36–38^.

PVDF-TrFE is a ferroelectric polymer that through extrusion processes can be formed into nanofiber scaffolds which mimic the intrinsic electrical and morphological properties of collagen type I (Fig. 1c). Mechanical actuation of these scaffolds through repetitive physiological loading and unloading (stretching) can deliver local bioelectrical cues to modulate tissue regeneration without the need for an external battery.

Here, we have developed a self-powered bioelectronic nanofibrous scaffold that serves to mechanically support the regeneration of damaged tendon tissues, maintain the tenogenic potential of endogenous tendon-derived stem cells and promote tendon regeneration, demonstrating that electromechanical stimulation can regulate the functional response and phenotypic maintenance of tendon cells. Additionally, we improved the physicomechanical and piezoelectric characteristics of the scaffold through near-field electrospinning intending to improve therapeutic performance. Finally, we established a rat model of tendinopathy and we demonstrated the efficacy of the implantable piezo-bioelectronic device in modulating the expression of ion channels (TRPA1, PIEZO2 and KCNK2/4) and activating specific signalling pathways (MAPK/ERK) associated with the endogenous tendon regeneration process.

### Design and fabrication method of the bioelectronic device

The biophysical properties of the bioelectronic scaffold, including its mechanical behaviour, microstructure (fibre diameter and alignment), and crystallinity-dependent piezoelectric response, play a pivotal role in regulating the cellular events involved in tissue regeneration^39^. To develop a collagen electrical-analogue device that can recapitulate the fibrous structure of tendon, an electrospinning process was pursued, and nanoscale fibres with varying diameter and alignment were produced through optimisation of solution and processing parameters (Fig. 2).

**Fig. 2.**
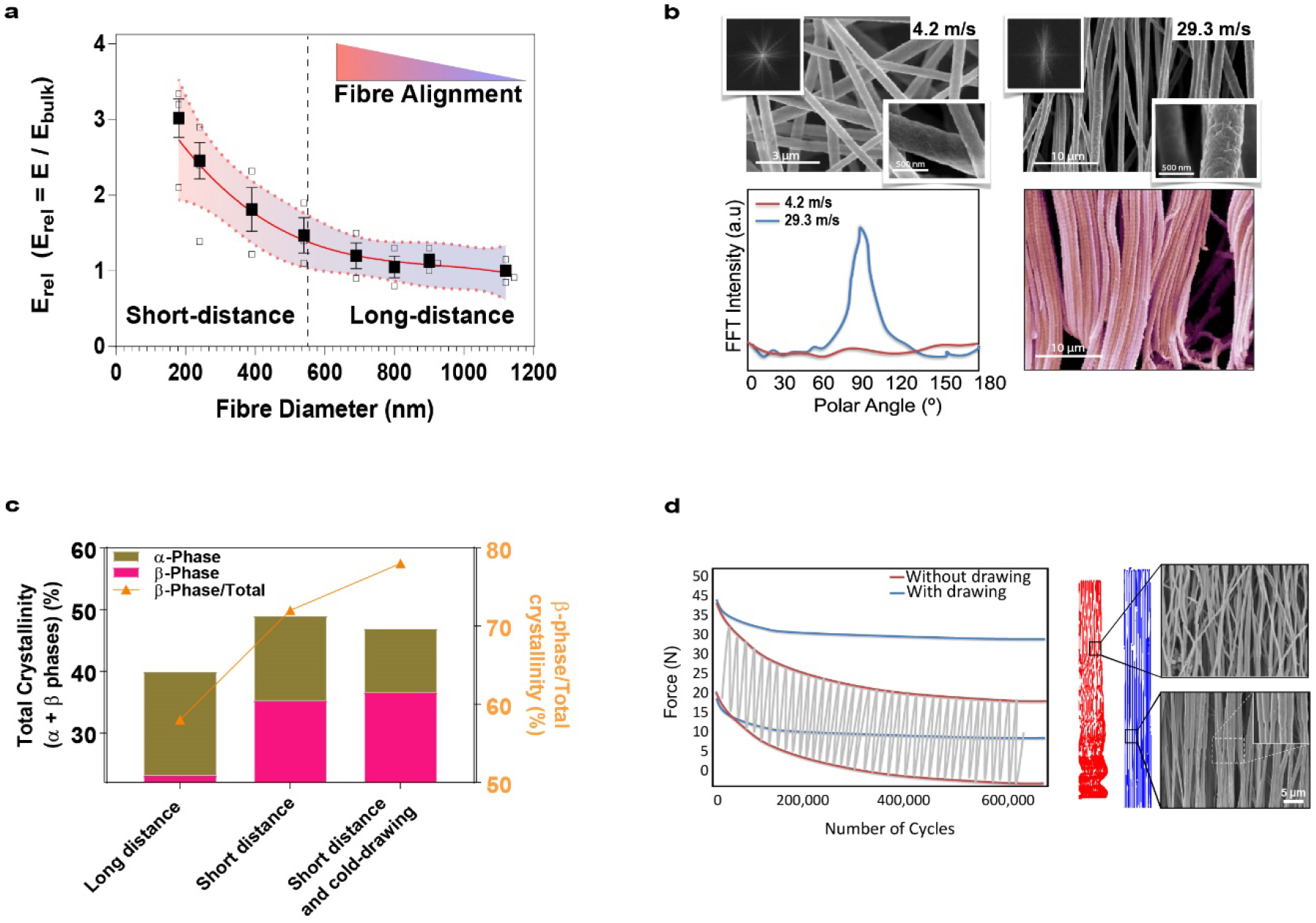
Cold drawing improves fibre alignment and enhances electroactive β-phase formation. **a**, Analysis of the relationship between fibre diameter and the relative Young’s modulus (E_rel_) indicated that the mechanical behaviour of highly aligned scaffolds was inversely proportional to fibre diameter **b**, The morphology of scaffolds obtained by electrospinning collected at low (4.2 m/s rpm) and high (29.3 m/s) linear speeds. Fast Fourier Transform (FFT) spectra showed a broad distribution of intensities for low speed (4.2 m/s) and a clear peak for high speed (29.3 m/s) characteristic of highly organised structures. Fibre alignment was significantly increased, and the morphology mimicked that of tendon collagen. **c**, Total crystallinity and β-phase content (calculated from FTIR and XRD spectra) increased as a function of processing parameters. **d**, Creep/stress relaxation was minimised after cold drawing and resulted in reduced fibre diameter and increased fibre alignment. SEM images of scaffolds with and without cold drawing.

First, we analysed the impact of fibre diameter on the biophysical properties of the scaffolds. By varying the concentration of the polymeric solution (1.1, 1.2, 1.3, 1.4, 1.5, 1.6 and 1.7 mg/ml in dimethylacetamide (DMAc) we could obtain fibres with distinct diameters. We used a 30 kV potential difference between collector and nozzle tip (200 μm), a flow rate of 1 ml/hr, and a tip-to-collector distance of 6 cm (short-distance electrospinning). Importantly, a collector disk with an 8 mm width was used to concentrate the electrical field while rotating at 29.3 m/s (linear speed) to draw and align the collected fibres. The resulting fibres ranged from 180 to 540 nm in diameter (Fig. 2a and Supplementary Fig. 1), which is similar to the diameter of collagen fibres in tendon tissue 50-500 nm^40^.

It was observed that the fibre diameter governed the stiffness of the resulting scaffolds and constructs with lower fibre diameters possessed an increased Young’s modulus, as determined by tensile-test analysis (Fig. 2a). The observed enhancement of the mechanical properties of scaffolds with decreasing fibre diameter was a result of a high level of chain extension and orientation along the fibre axis as a consequence of fibres being drawn by centripetal forces during fibre collection. With semi-crystalline polymers (including PVDF-TrFE), it has been demonstrated that stress hardening does not depend on polymer crystallinity but rather, on the density of amorphous polymer chain entanglement, which in respect to electrospinning, is related to the distance between the needle and collector, the collector rotational speed and the polymer solution concentration^41–43^. Finally, nanofibers with diameters <240 nm (Supplementary Fig. 1f) were prone to alignment disorganisation due to the air currents generated during collection. As a result, scaffolds with a fibre diameter between 390 and 540 nm (Supplementary Fig. 1d,e) demonstrated increased fibre alignment.

To decouple the effect of piezoelectric stimulation from mechanical loading on the cellular response, a non-piezoelectric PTFE scaffold which maintained the topography (i.e. fibre diameter and organisation) and was chemically analogous (i.e. a fluorinated polymer) to PVDF-TrFE was also fabricated. PTFE is non-piezoelectric due to the presence of a centrosymmetric structure with strong C-F polar covalent bonds (C∂+–F∂−). Conversely, PVDF-TrFE presents non-centrosymmetric electrical dipoles, generated by the high electronegative difference between hydrogen and fluorine atoms (Fig. S2). The direct electrospinning of PTFE fibres, however, was not possible due to the unique chemical properties of PTFE. Therefore, a two-step near-field electrospinning process was optimised to obtain scaffolds of PTFE fibres with a diameter of 540 nm (Supplementary Fig. 2)^44^.

The tendon microstructure is highly hierarchical, with collagen fibres aligning to the direction of mechanical loading. To reproduce this critical feature in both PVDF-TrFE and control PTFE scaffolds the collector velocity was adjusted between 4.2 – 29.3 m/s, and the concentration of the polymeric solution fixed at 1.7 mg/ml. As evident in Fig. 2b, the scaffolds obtained at a collector linear velocity of 29.3 m/s displayed a highly organised fibre morphology, while a broad distribution of fibre orientation was observed for scaffolds formulated with a collector linear velocity of 4.2 m/s. Additionally, as determined by X-ray diffractometry (Supplementary Fig. 3), the degree of crystallinity increased from 40% in randomly aligned PVDF-TrFE fibres to 49% in aligned fibres (Fig. 2c), suggesting that mechanical drawing during fibre collection at high rotational speeds resulted in PVDF-TrFE fibres with a reduced diameter (590±130 nm for 4.2 m/s and 540±120 nm for 29.3 m/s) and enhanced crystallinity (Supplementary Table 1).

Our near-field electrospinning configuration contrasted to previously reported electrospinning configurations for PVDF-TrFE (i.e. long-distance electrospinning)^45^. We have demonstrated that scaffolds produced using a near-field configuration possessed increased fibre organisation (Supplementary Fig. 1) and crystallinity (Fig. 2c), supporting the hypothesis that fibres are significantly drawn when subjected to an electric field of increased intensity. The mechanical behaviour of the scaffolds obtained by the short-distance configuration was improved to their long-distance counterparts in terms of elastic modulus, creep resistance and yield strength (Fig. S4), which is of vital importance for the device to retain the integrity of its macro and microstructure following implantation^46^. Regarding the piezoelectric response, a growing body of research suggests that the piezoelectric performance of PVDF (i.e. the base comonomer of the PVDF-TrFE copolymer) is affected by the degree crystallinity (Fig. SI4 and Supplementary Table 2) and in particular the β-phase content^47,48^. As determined by Fourier-transform infrared spectroscopy (FTIR) (Supplementary Table 3), the short-distance configuration promoted the crystallisation of the electroactive β-phase (Fig. 2c). Overall, these observations reinforce the argument for the potential use of near-field electrospinning in the fabrication of PVDF-TrFE-based scaffolds.

To further increase the mechanical properties and β-phase content of the scaffolds, while avoiding the well-reported crimping of the fibres under repetitive mechanical loading, two post-synthesis processes were investigated: i) strain-hardening (cold-drawing) and ii) thermal annealing (cold crystallisation). Based on the initial properties of the scaffolds, we explored cold drawing at ~12% to induce strain-hardening through plastic deformation of the structures. Overall, this process resulted in ~8% plastic deformation, alignment of fibres (from 65% to 98%) and reduction in individual fibre diameter (from 540 to 516 nm). Importantly, fibre diameter reduction and better fibre alignment resulted in scaffolds with an improved elastic modulus (61.8 ± 8.1 MPa) and strength (31 ± 4.2 MPa) (Table 1).

**Table 1.** The physical properties of piezoelectric PVDF-TrFE and non-piezoelectric PTFE scaffolds

| Scaffold Type | Elastic Modulus (MPa) | Strength (MPa) | Elongation (%) | Fibre diameter (nm) | d<sub>33</sub> (pC/N) |
| --- | --- | --- | --- | --- | --- |
| Non-piezoelectric with drawing | 14.5 ± 1.7 | 16 ± 0.3 | 91.3 ± 10.4 | 690 ± 110 | 0 |
| Piezoelectric w/o drawing | 56.6 ± 7.6 | 15 ± 3.7 | 46.4 ± 6.1 | 540 ± 120 | 29.3 ± 2.7 |
| Piezoelectric with drawing | 61.8 ± 8.1 | 31 ± 4.2 | 39.4 ± 3.2 | 513 ± 80 | 36.5 ± 3.9 |

Concurrently, excess stress relaxation/creep of the fibres was significantly reduced (Fig. 2d). Thermal-annealing around the T_c_ (90°C for 1 hour) was used to further increase the β-phase content. The amount of β-phase content in PDVF-TrFE samples subjected to post-synthesis cold-drawing and thermal annealing was higher than the non-post-treated counterparts as determined by FTIR analysis (Fig. 2c).

The effect of alignment on the piezoelectric performance of individual fibres was demonstrated by switch-spectroscopy piezoresponse microscopy (SS-PFM). The piezoelectric response of fibres from randomly-aligned scaffolds was compared to that of fibres from mechanically drawn, aligned scaffolds. The piezoresponse or d_33_ value was found to be dependent on the fibre alignment. A strong correlation was observed between scaffolds with increased fibre alignment and a higher piezoelectric response (from −16.92 to −24.61 pm/V) (Fig. 3). The elastic modulus of individual fibres was measured using peak-force imaging. The apparent modulus of individual fibres was 350 ± 80 MPa, yet no significant differences were observed between random and aligned fibres. Interestingly, a cooperative piezoelectric effect previously described by Persano *et al*.^49^ was found in dense arrays of fibres (Fig. 3a) due to electromechanical interactions between adjacent fibres and the scaffold stiffness gradients, resulting in an overall enhancement of the scaffold piezoresponse (d_33_ = −36.5 ± 3.8 pm/V and g_33_ = −0.41 V.m/N). Differences between single fibres and multiple fibres (arrays) arise from local differences in stress distribution variations. Discrepancies between obtained values and reported values (d_33_ = −29 pC/N) may arise from differences in fibre diameter, orientation, geometry and arrangement^50–54^.

**Fig. 3.**
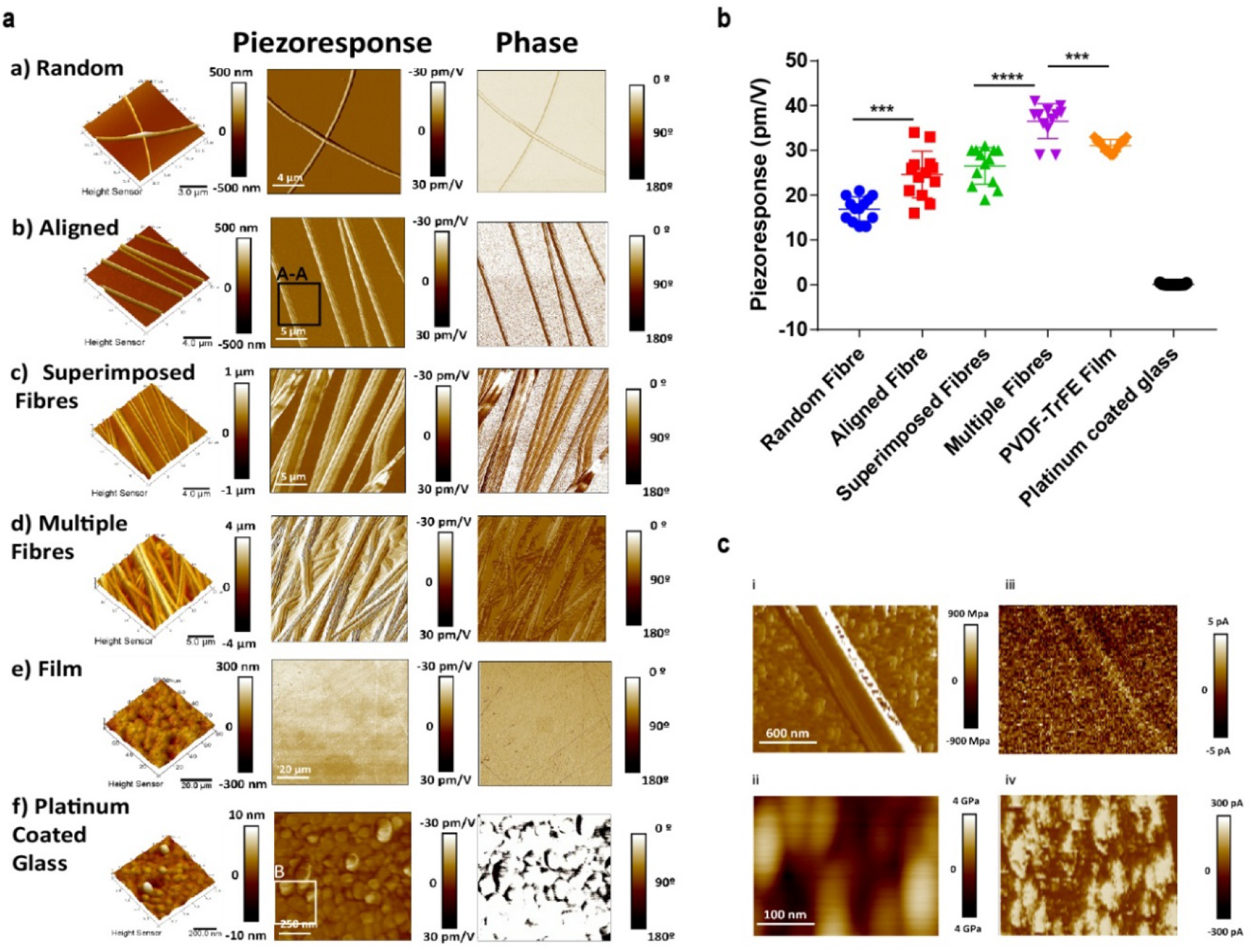
Fibrous aligned piezoelectric scaffolds demonstrated higher piezoelectrical performance than piezoelectric films due to geometrical boundary variables that control the piezoresponse of individual PVDF-TrFE fibres. PFM amplitude and phase images for single and multiple fibres. **a**, Random individual fibres (a) demonstrated lower piezoelectric performance relative to aligned individual fibres (b). Aligned superimposed (c) displayed an enhanced piezoelectric response (in-plane), showing a cooperative effect due to the electromechanical interaction among adjacent fibres. Multiple dense layers (d) of adjacent fibres presented the highest piezoelectric coefficient. Commercial PVDF-TrFE films (**e**) display a lower piezoelectric coefficient. A non-piezoelectric platinum substrate showed no piezoresponse (f). **b,** Direct comparison of d_33_ values (piezoresponse) between samples (a-f). **c. i** Individual fibre DMT Elastic modulus and **iii** electrical conductivity. **ii** Electrical currents and **iv** DMT elastic modulus were measured on a platinum-coated glass surface as control, I = 300 pA The relative stiffness difference between single fibre (350 ± 80 MPa) and fibre arrays is responsible for the out-of-plane piezoresponse enhancement. No residual (I = 0 pA) electrical currents were measured in the individual fibres indicating no resistive mode of conduction characteristic of piezoelectric materials (All measurements were obtained using N=3 samples, r=7 replicates per sample). The values are presented by mean ± SD. Significant differences (one-way ANOVA) of piezoporesponse of fibres. Asterisks (*p<0.05, **p<0.001) indicate results of post-hoc test (Bonferroni).

Mechanical and electromechanical stimulation of cells was performed *in vitro* using non-piezoelectric PTFE and piezoelectric PVDF-TrFE scaffolds under cyclic mechanical stretching. The physical and electrical properties of the scaffolds are shown in Table 1.

### The role of electromechanical stimulation on tendon-specific gene expression and phenotypic maintenance *in vitro*

Tendons have a great ability to respond to mechanical forces by adapting their physicomechanical structure and biochemical composition. Cyclic mechanical stretching (progressive rehabilitation) of the tendon is an excellent and recognised method for treating tendon-related injuries and benefits overall tendon health^55–57^. Several biomechanical studies have confirmed that tendon is subjected to 3-4 % strain during normal activities^58–61^ and also, that strain rates (loading frequency) are important in modulating the cellular response *in vitro*^62–64^. Under physiological conditions, the strain rate of the tendon is around 0.1 – 0.5 Hz; however, during intense activity, the frequency can be as high as 10 Hz^65,66^. Interestingly, piezoelectric systems exhibit frequency-modulated performance (output charge) due to the frequency dependence of the elastic/piezoelectric constants. The effect of strain rate/frequency (0.25 – 3 Hz) was evaluated using physiologically relevant magnitudes of 1%, 2% and 4% strain (Fig. 4c). Aligned PVDF-TrFE scaffolds produced a significant piezoelectric response with the application of force parallel to the fibre orientation (Fig 4a, c). The results showed a positive correlation between voltage output and strain magnitude demonstrating the piezoelectric effect of the scaffolds. As shown in Fig. 4b, the piezoelectric behaviour of the scaffolds was further confirmed by using SS-PFM.

**Fig. 4.**
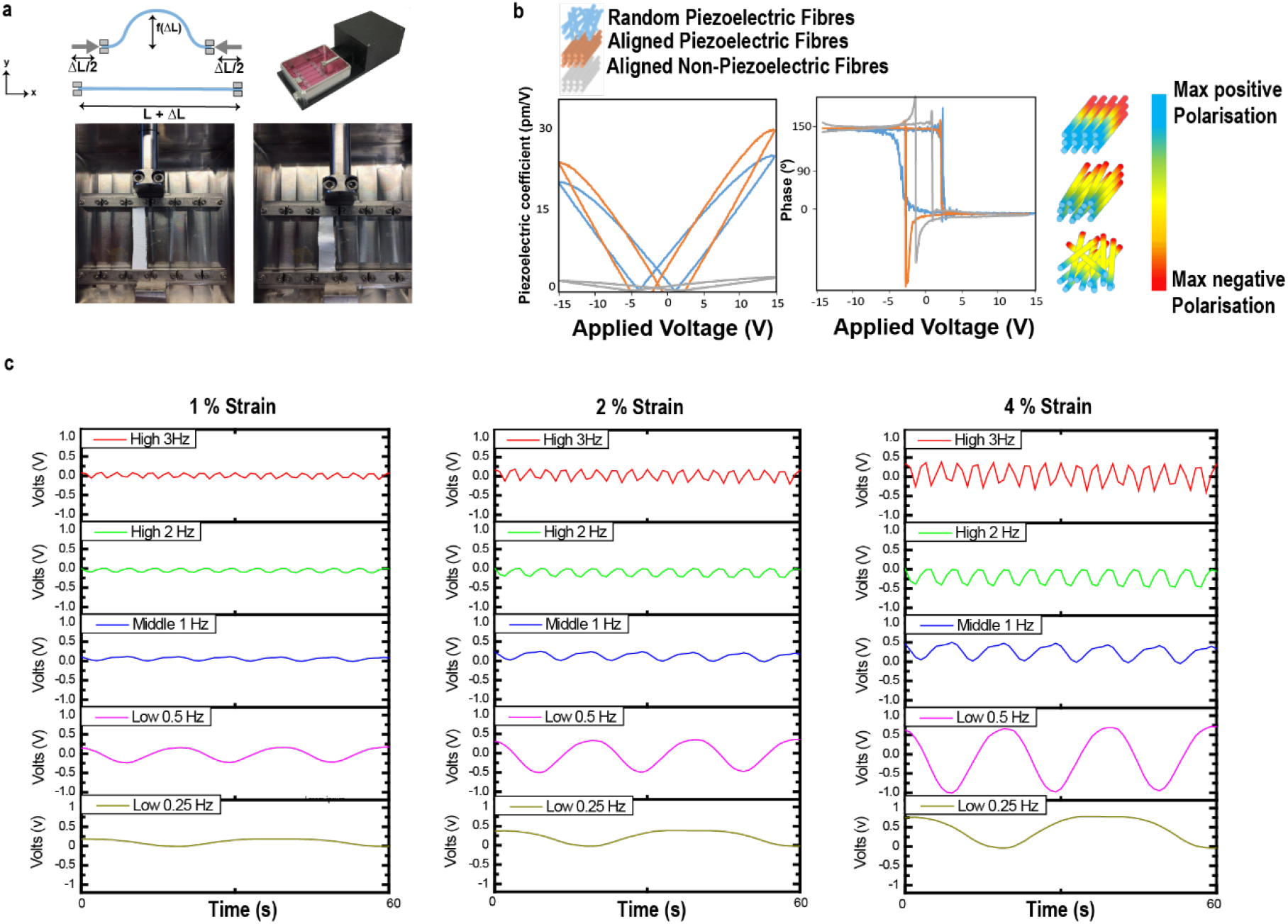
The electrical output of PVDF-TrFE scaffolds varies as a function of frequency/strain rate and strain amplitude. **a**. An overview of the mechanical loading system used to measure voltage output and used for the *in vitro* studies. **b**. Representative hysteresis and butterfly loops of the fibre piezoresponse and polarisation. SS-PFM facilitated the measurement of the PFM phase and amplitude response loops as a function of the applied voltage. The amplitude image shows the magnitude of displacement response generated under a controlled applied voltage in drawn fibres, and the highest amplitude peak was shown to be around 300 pm under 12 V. The phase image shows the positive and negative values of antiparallel ferroelectric nanodomains. The phase measurements indicated a ~180° switch of the dipoles between the applied positive and negative voltages. The piezoresponse obtained from PFM measurements showed that the electrical dipoles lie normal to the surface, characteristic of the organised electrical dipoles/chains of β-phase crystals. PTFE did not demonstrate piezoelectric behaviour (grey line). The schematic indicates the voltage distribution on the fibre surface as a function of geometrical arrangement (fibre orientation) and interaction with adjacent fibres that result in differing transverse deformation restriction and modulation of piezoresponse along the longitudinal fibre axis. **c**. Voltage measurements as a function of frequency (0.25, 0.5, 1, 2 and 3 Hz) at a constant amplitude of 1%, 2% and 4% strain.).

The scaffold bias voltage output was observed to increase proportionally with increasing strain from 0.28 to 1.21 V at a fixed frequency of 0.25 Hz (1-4 % strain). With a fixed strain amplitude of 4%, the range of voltage output ranged from 0.51 V (3 Hz) to 1.21 V (0.25 Hz) (Fig. 4c). Furthermore, stability analysis of up to 500 cycles of continuous dynamic strain did not reveal any significant changes to this output (Fig. 2d).

It has widely been demonstrated that tendon cells are sensitive to mechanical loading, which can lead to modulated gene expression, protein synthesis and mitogenesis *in vitro* through activation of mechanotransductive processes^11,17,67–71^. Here the role of electromechanical stimulation on tenocyte function was evaluated *in vitro.* Following scaffold fabrication, the fibres were covalently functionalised with fibronectin to enhance cell adhesion and to sustain cell proliferation (Supplementary Fig. 5)^72^. First, the sole effect of topographical cues present on the different scaffolds and films, on cell morphology and organization and, tendon specific markers Tenomodulin (TNMD) and Scleraxis (SCX), was evaluated by seeding human tenocytes onto aligned, random (non-aligned) scaffolds and planar films for one, three and seven days under static conditions (Fig. 5a,c). The transcription factor Scleraxis (SCX) is highly specific for both mature and precursor tendon cells and is considered the master regulator of the tenocyte phenotype and tendon-specific differentiation^73^. Similarly, Tenomodulin (TNMD) is a glycoprotein involved in tenocyte proliferation and is an established marker for tenocyte differentiation. Our observations confirmed that the induced elongation by contact guidance of cells on aligned scaffolds resulted in relative SCX and TNMD upregulation relative to planar films and non-aligned scaffolds (Fig. 5 a,c and Supplementary Fig. 6). Following optimisation of mechanical loading condition through analysis of tenocyte viability, nuclear deformation and TNMD and SCX expression (Fig. 5b and Supplementary Fig. 6), 0.5 Hz frequency was chosen due to comparable cell viability compared to static conditions (Fig. 5b). Interestingly, the expression of TNMD was significantly higher than the static control after five and day ten days in culture with a fold-increase of ~1.5 and ~2, accordingly (Fig. 5d).

**Fig. 5.**
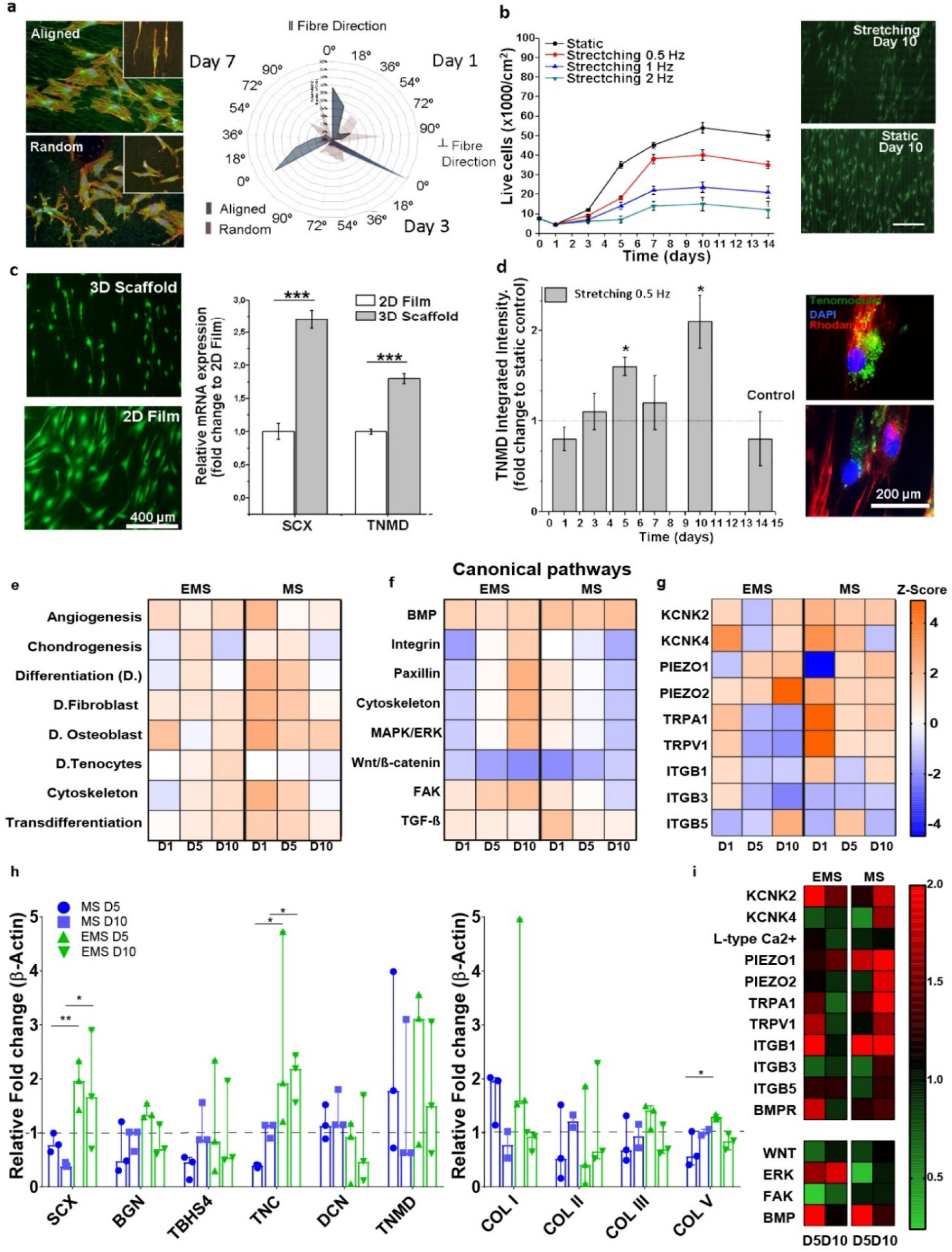
Electromechanical stimulation induces maintenance of tendon specific gene expression and phenotype in vitro. **a**, Human tenocytes demonstrated an elongated morphology from day 1 when cultured on aligned PVDF-TrFE scaffolds. Inserts are higher magnification (red = actin, green = vinculin, and blue = nucleus, scale bar = 400 um). Angle distribution radar analysis indicated that the percentage of aligned cells was significantly modulated by fibre direction at day 1, 3 and 7 (0° and 90° correspond to alignment and no alignment respectively). **b**, Tenocytes cultured on piezoelectric scaffolds under dynamic stimulation subjected to different strain rates (0.5, 1 and 2 Hz) exhibited proliferation rates inversely proportional to the strain frequency. **c**, Cells demonstrated differential expression of tenospecific proteins and morphological changes when cultured on electrospun scaffolds or planar PVDF-TrFE films. Tenocytes maintained their phenotype and expression levels of TNMD and SCX when cultured on aligned piezoelectric scaffolds; this effect was not present with 2D planar films. **d**, An increase in the expression of TNMD protein expression was observed in cells subjected to 0.5 Hz electromechanical stimulation at days five and ten. **e**, Functional analysis of human tenocytes cultured under mechanical and electromechanical stimulation for one, five and ten days. The significant changes in gene expression were classified into broad signalling pathways (**e**), specific biological processes (**f**) and ion channels and integrins (**g**). In particular, electromechanical stimulation for ten days induced a significant activation in tendon specific differentiation pathways associated with activation of MAPK signalling pathways and activitydependent downregulation of TRPA1 and Piezo2 ion channels. This trend was significantly reversed in cells exposed to mechanical stimulation alone (Median ± ra* p<0.1, ** p<0.01). **h**, Expression profiles of tendon specific proteins and collagens in tenocytes subjected to electromechanical (EMS) and mechanical (MS) stimulation at day five (D5) and day ten (D10). A significant increase of Col. V synthesis is observed during EMS relative to MS at day five. **i**, Heatmap of ion channel, integrin and tendon/osteochondral specific protein expression in tenocytes subjected to EMS and MS at day five and ten (Median ± range, N=3).

Subsequently, genomic analysis of human tendon-derived cells cultured on aligned and drawn piezoelectric PVDF-TrFE and non-piezoelectric PTFE scaffolds under both static (0 Hz) and dynamic conditions (0.5 Hz, 4% strain) was performed for one, five and ten days. Overall, the analysis of gene expression correlated well with the results observed at the protein level (Fig. 5d). Electromechanical stimulation (mechanically stretched PVDF-TrFE relative to static PVDF-TrFE) induced a rapid and sustained up-regulation of tendon-related and bone-related genes. Conversely, mechanical stimulation (mechanically stretched PTFE relative to static PTFE) induced a significant up-regulation of bone and tendon-related genes at day one, although this was significantly decreased at day five and ten (Supplementary Fig. 7a). By day ten, bone-related genes were up-regulated in tenocytes subjected to both mechanical and electromechanical stimulation (Supplementary Fig. 7a). Interestingly, at day ten SCX and TNMD were upregulated only in the electromechanical stimulation group (Fig. 5e). Taken together, these observations indicate that trans-differentiation of tenocytes towards an osteogenic/chondrogenic lineage can be modulated with mechanically loaded piezoelectric (PVDF-TrFE) and non-piezoelectric scaffolds (PTFE). Moreover, electromechanical stimulation offered sustained tenogenic differentiation capacity relative to mechanical stimulation alone.

To unravel the signal transduction pathways responsible for the regulation of tenogenic differentiation, ingenuity pathway analyses (IPA) was used with gene expression profiles. IPA attributes an activation z-score used to predict the activation state of biological functions and signalling pathways. Pathway analysis of human tenocytes yielded significant alterations of broad functional signalling pathways, and a mechanistic network of genes and specific biological functions were differentially regulated by both electrical and mechanical stimulation (Supplementary Fig. 8). Canonical signalling pathways with the greatest number of modulated gene expression were associated with mechanical stimulation at day one and electromechanical stimulation at day ten (Supplementary Fig. 7b). TGF-β/BMP signalling was consistently activated through both types of stimulations; however, it was significantly increased in tenocytes subjected to mechanical loading at days one and ten (Fig. 5e,f). Similarly, increased activation of osteospecific cell differentiation and Wnt/β-Catenin signalling pathways were attributed to mechanical activation at day five and ten relative to EMS. Importantly, following ten days in culture, the tendon-related signalling pathway MAPK/ERK was activated in tenocytes subjected to electromechanical stimulation (PVDF-TrFE) but downregulated under continuous mechanical stimulation (PTFE) (Fig. 5f).

During tendon specific differentiation or trans-differentiation, an orchestrated signal transduction events take place leading to the temporally regulated expression of genes associated with the tendon-specific phenotype or phenotypic drift respectively. We hypothesized that mechanically activated membrane structures and electrically gated ion channels would undergo modulated expression in response to electromechanical stimulation. To investigate the role of these genes on signal generation and propagation, correlation analysis was conducted between the obtained significant gene modulations at day one, five and ten in the mechanistic network with differential changes in the expression of mechano-sensitive (TRP family ion channels, focal adhesion-related proteins and integrins), electro-sensitive (KCNK family and Ca^2+^ L-type ion channel), and piezo-sensitive (Piezo family ion channels) receptors. Critically, increased activation of tendon differentiation-related genes (Fig. 5e) through the modulated activity of tendon-related transcription factors (i.e. SCX and EGR1, see Supplementary Fig. 7) was associated with a decrease in the expression level of TRP family ion channels under electromechanical stimulation at day 5 (Fig. 5g). As shown in Fig. 5h, SCX and TNC were significantly upregulated (see Fig. 5) under electromechanical stimulation at day 5 and 10 compare to mechanical stimulation and associated with down-regulation of PIEZO2, TRPA1 and TRPV1 ion channels genes(Fig. 5g,i).

Conversely under mechanical stimulation only, a down-regulation in the expression of tendon-related genes and up-regulation in expression of genes associated with cartilage/bone differentiation was associated with an up-regulation of TRP ion channels (Fig. 5e and Supplementary Fig. 8). Specifically, TRPA1 and TRPV1 ion channels underwent sustained up-regulation and correlated with the positive regulation of transcription factors SOX9, HDAC1 and SP7, proteins that have a recognised role in the process of osteogenesis (Supplementary Fig. 7).

Electromechanical stimulation also resulted in the upregulation of several genes involved in bone differentiation. For instance, multiple bone transcription factors such as RUNX2 and BMP2 were activated, indicating that the electromechanical stimulation, can lead to coactivation of both bone and tendon related signalling pathways in human tenocyte populations. These results were further validated using a protein array demonstrating that mechanical stimulation resulted in significant upregulations of ion channels (KCNK4, PIEZO1, PIEZO2, TRPA1, TRPV1) whereas electromechanical stimulation leads to a downregulation of almost all ion channels correlated to the activation of the MAPK/ERK pathway (pERK/tERK, Fig. 5f,i).

### Electromechanical stimulation induces the activation of tendon specific signalling pathways in vivo

Over the last century, a clear link has been established between biomechanical loading and tissue function. Although significant progress has been made in the field of cellular mechanobiology, the exact mechanism of how cells can sense and respond biochemically to mechanical and electrical forces still remains elusive^74–76^. An increasing body of research indicates that cells utilise different molecular machinery for sensing distinct external stimuli with high spatiotemporal resolution^74,77^. The biological response can range from cytoskeletal reorganisation, modulated ECM adhesion or cellular reprogramming^78^. At the tissue level, transient mechanical cues result in altered cellular function and subsequent changes in the composition of the interstitial matrix, reaching a homeostatic balance between mechanical loading and tissue mechanics^14,79,80^. Therefore, to investigate the effect of electromechanical stimulation on tendon repair through our piezoelectric scaffold *in vivo,* the development of a preclinical load-bearing injury model was necessary.

We, therefore, developed a full-thickness Rat Achilles tendon defect model (Fig. 6a and Supplementary S9a) to investigate tendon repair and tendinopathy progression processes in response to electromechanical stimulation, where rats were subjected to either a daily treadmill running regime (TR) or to cage roaming conditions only (Cage) (Fig. 6a). To explore in detail the recovery of function, the animal’s gait was analysed during treadmill exercise which suggested that treadmill running had a consistent significant positive effect in animal motility following a tendon injury. In particular, at week eight, both experimental groups which received the non-piezoelectric and piezoelectric scaffolds (TR) displayed a significant increased of their functional recovery in comparison to static controls (Cage) (p<0.05 and p<0.05 respectively, Fig. 6a,e and Supplementary Table 4).

**Fig. 6.**
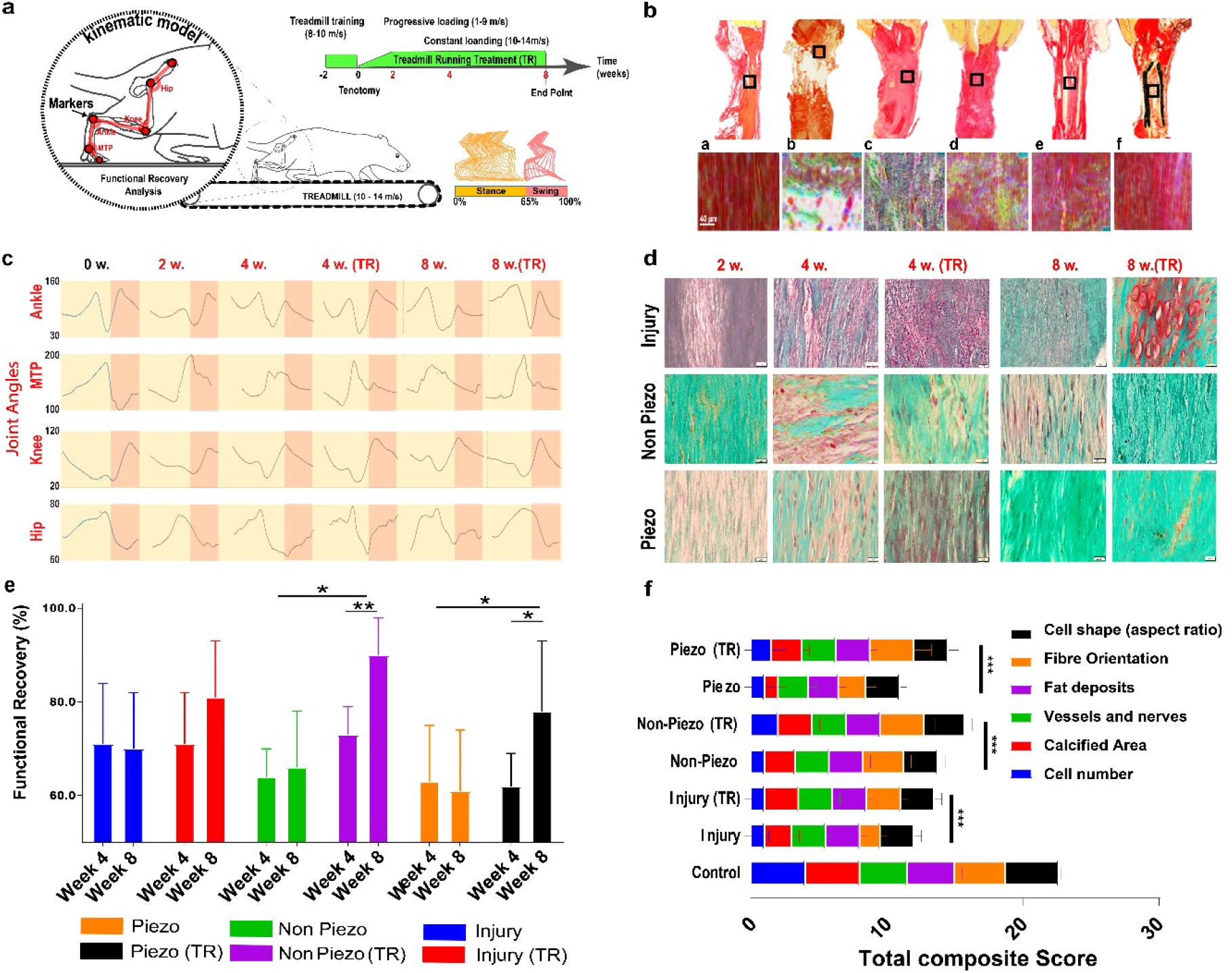
Tendon regeneration and tissue organisation are enhanced in animals treated with mechanical and electromechanical stimulation in a tendon defect model. **a,** Schematic representation and experimental timeline of the Achilles tendon defect repair model and kinematic model for gait analyses. **b**, Picrosirius red staining of intact, suture-repaired and scaffold-repaired tendon tissue. Orientation distribution analysis using high-magnification images (box region) indicated that tissue organisation was increased in animals treated with piezo and non-piezo scaffolds. a) intact tendon, b) sutured tendon 24 hours post-injury, c) 1-week post-surgery and d) 8 weeks post-injury; e) tendon repaired with piezoelectric scaffolds 8 weeks post-injury and f) tendon repaired with a non-piezoelectric scaffold 8 weeks post-injury. **c**, Gait analysis for all groups was obtained through analysis of four different joint angles (Ankle, MTP, Knee and Hip) at weeks 2, 4 and 8 post-injury. The angle amplitudes obtained for animals treated with piezoelectric scaffolds after surgery (injury) and subjected to treadmill running increased over 8 weeks—example of a still image obtained from video recordings used for the measurement of gait. **d**, Representative Safranin O stained histological images indicating the role of scaffold medicated mechanical and electromechanical stimulation on proteoglycan and collagen II deposition and chondrocyte distribution in animals subjected to the treadmill and static conditions. Tendon defects repaired with non-piezo and piezo scaffolds and subjected to treadmill loading demonstrated less intense proteoglycan (GAG) (red) and collagen II (green) staining relative tendon tissues retrieved from the cage groups. Suture repaired tendon tissues possessed a highly disorganised structure with increased GAG and collagen II content and increased chondrocyte presence. **e**, Composite functional recovery analysis. It was obtained a significant increase in both MS and EMS (TR compared to Cage) at week eight. **f**, Total composite score of the tissue quality at week eight. It was obtained that all TR groups showed significant increase with respect to cage. Scale bar is 200 μm.

Under non-pathological conditions (non-sectioned tendons) cells showed an elongated phenotype (Supplementary Fig. 9b,c) and resided in a rich and dense ECM composed of predominantly collagen type I cross-linked by GAGs (Supplementary Fig. 10,11). In our Achilles-tendon injury model resident and invading cells adopted a non-elongated, non-aligned morphology and after eight weeks post-injury, developed calcific tendinopathy (Supplementary Fig. 10). Moreover, micro CT scanning in animals treated with clinical standard suture stabilisation (injury), confirmed the presence of ectopic bone tissue at eight weeks post-injury (Supplementary Fig. 12, 13).

Significant increases in tissue organisation (Fig. 6f and Supplementary Table 4) were observed in tendon tissues derived from animals which were treated with mechanical (non-piezoelectric PTFE scaffold and treadmill running) or electromechanical stimulation (piezoelectric scaffold and treadmill running) following tendon injury (Fig. 6b). Scaffold-mediated tendon repair reduced ectopic bone and enhanced mature tendon tissue formation (Fig. 6b). Furthermore, it was observed that loading conditions (Piezo TR, Non-piezo TR) using treadmill running resulted in a rapid regain of animal motility and functional recovery (Fig. 6c and e), which was associated with a significant decrease (p<0.05) in the presence of cartilage-related glycosaminoglycans (GAGs), as quantified via Safranin-O staining intensity (Fig. 6d and Supplementary Fig. 13). Furthermore, an increase in the composite tendon tissue regeneration score was obtained with the use of non-piezoelectric scaffolds relative to the piezoelectric scaffold in mechanically stimulated animals (TR) (Fig. 6f and Supplementary Table 5).

Custom array analysis of proteins extracted from tendon tissue obtained from all experimental and control groups indicated differential expression of tendinopathy associated protein collagen II and tendon specific proteins (Fig. 7). Generally, collagen II is only expressed in tendon tissue at the regions of bone insertion; interestingly, it was observed that under cage roaming conditions, tendon injuries treated with the piezoelectric scaffold exhibited a relatively significant increase in calcification eight weeks post-injury compared to the other groups (Supplementary Fig. 12). Critically, collagen II expression was not increased under electromechanical stimulation (Piezo and TR) relative to mechanical stimulation, demonstrating the regulatory potential of dynamic electrical cues on tissue composition (Fig. 7a).

**Fig. 7.**
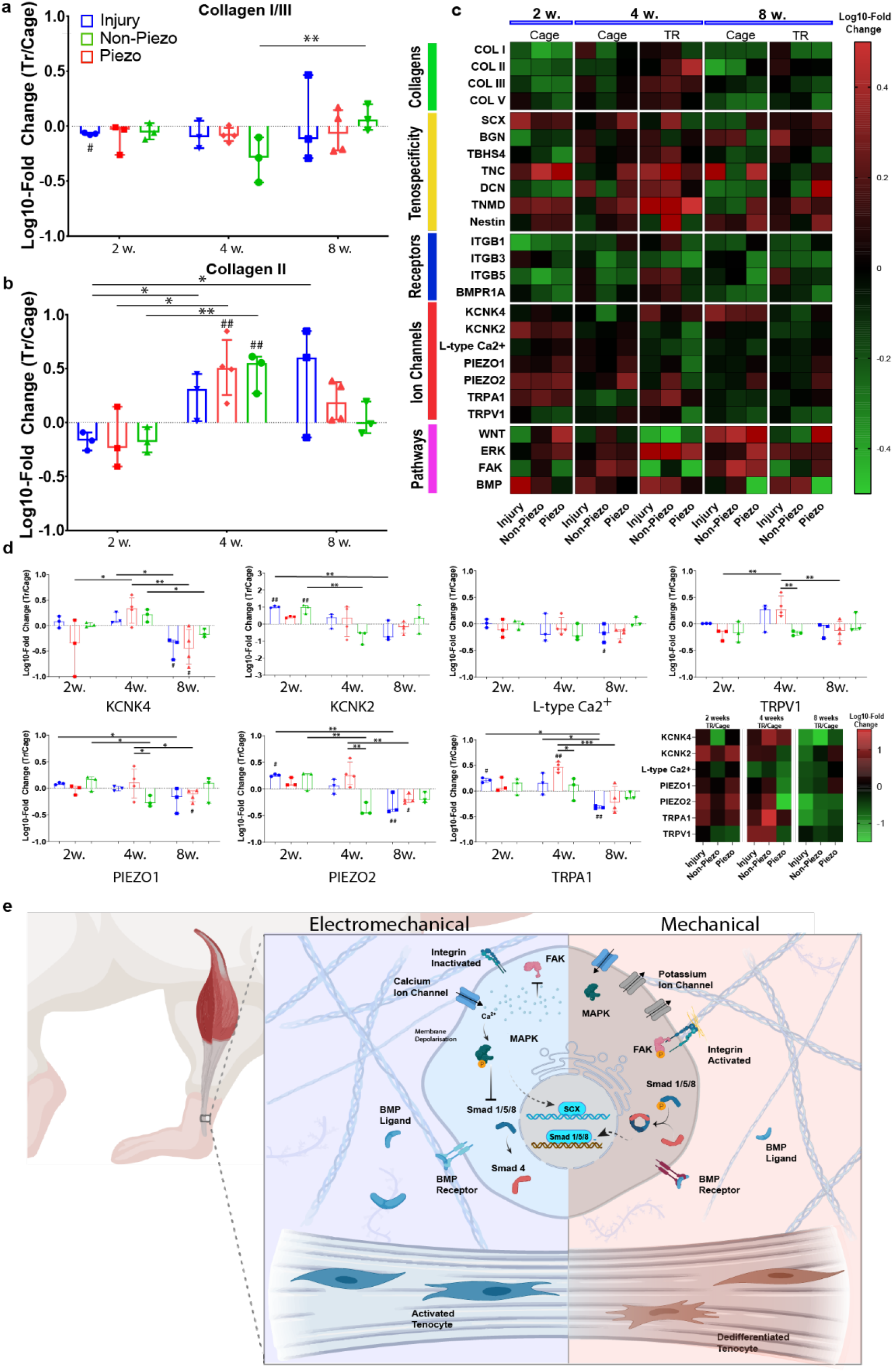
Electromechanical stimulation promotes collagen synthesis and controls tendon-specific protein expression. **a,** Collagen II expression comparison between animals subjected to treadmill running (TR) and cage conditions (Cage) at 2,4 and 8 weeks (results are represented as mean ± SEM). **b** Collagen expression log10-fold change and p-values (animals subjected to treadmill running vs. animals in caged conditions, TR/Cage). (* p<0.05;** p<0.01; *** p<0.005 with respect to same Group at 2, 4 and 8 weeks, # p<0.05; ## p<0.01 with respect to control Cage at same time point). **c**, Heatmap of protein expression log10-fold change (treated healed tissue vs contralateral tendon tissue) The effect of mechanical and electromechanical stimulation on the expression of proteins associated with cellular receptors, pathway functions, ECM production and molecular mechanisms related to tenogenesis was assessed via protein microarray. Heat maps of proteins analysed across samples collected from injury, non-piezo and piezo groups at weeks 2, 4 and 8 (static and TR). The 35 proteins are represented in rows and classified in different groups according to their function. Data represents mean value of fold change relative first to β-actin and then to control sample (contralateral intact tendon) (N=4). Averaged intact tendon sample is represented by fold change mean value relative to β-actin of all contralateral samples collected at weeks 2, 4 and 8 for static and TR groups. Data are represented as mean ± SEM (n=3). One-way ANOVA with Tukey post hoc test was performed. *, **, *** Represents a statically significant difference (p<0.05), (p<0.01) and (p<0.001), respectively. **d** Ion channel expression log10-fold change and p-values (animals subjected to treadmill running vs. animals in caged conditions, TR/Cage). Significant altered expression of ion channels was detected for animals under MS or EMS. Significance threshold (* p<0.05;** p<0.01; *** p<0.005 with respect to same Group at 2, 4 and 8 weeks, # p<0.05; ## p<0.01 with respect to control Cage at same time point) **e**, The effect of electromechanical stimulation on mechanosensitive protein receptors and ion channel activation, and the suggested mechanism. Under normal physiological conditions, the activities of potassium channels limit membrane depolarization of tendon cells. In response to injury or mechanical loading, tyrosine kinase activity (i.e.FAK) regulates the expression of K and Ca channels at the transcriptional level although the transcription factors involved in this effect are unknown. Finally, a reduction in the expression of potassium channels results in increased membrane depolarization and calcium activity under the electrical stimulus.

Following injury, the synthesis ratio of collagen I/III was significantly increased in response to EMS at week eight (remodelling phase) with respect to week four (proliferative phase) to provide elasticity and mechanical support during defect stabilisation (Fig 7b). Furthermore, analysis of tendon-specific proteins SCX, Tenascin-C, Byglican, Tenomodulin, and Thrombospondin-4 was performed to evaluate tendon specificity during the repair. Protein array analysis revealed that SCX and TNMD were significantly upregulated at two weeks after tenotomy for all treatments and were differentially regulated after four and eight weeks of continuous mechanical or electromechanical stimulation. SCX and TNMD demonstrated significant upregulations (>2-fold change) during tendon healing (after eight weeks) in tendons treated with piezoelectric scaffolds (Fig. 7c). Furthermore, significant modulation of the tendon-specific functional response was observed at four weeks with respect to collagen synthesis and tenospecific associated proteins.

Several studies have shown that mechanical loading supports the maintenance of a tendon phenotype and sustained tendon-related protein synthesis^11,81^ through the activation of FAK signalling^82,83^. Conversely, excessive mechanical loading has been shown to induce tendon cell dedifferentiation towards an osteochondrogenic lineage via modulation of osteospecific BMP and Wnt signalling pathways^17,84^. Here, after continuous EM stimulation for eight weeks was observed to downregulate BMP signalling and upregulates the expression of tendon functional proteins including SCX and BGN (Fig. 7c). Additionally, upregulation of TNMD and downregulation of BMP signalling was associated with piezoelectric scaffolds in both static (cage) and dynamic (TR) conditions at four and eight weeks. To explore further the mechanism of the tissue response to mechanical or electromechanical stimulation, we investigated the correlation between the activation of mechanotransductive (MAPK and FAK) and osteospecific Wnt and BMP signalling pathways with of two different groups of proteins that act as “active sensors” of mechanical and electrical cues. The two groups were receptors (integrins β1, β3 and β5 and BMP receptor)^85–87^ and ion channels sensitive to mechanical and/or electrical stimuli (KCNK, Ca^2+^ L-type, TRPA1, TRPV1, Piezo1 and Piezo2) ^88–91^.

It was revealed that mechanical loading (TR) of tendon tissues for all treatments (Injury, Non-Piezo and Piezo) resulted in the upregulation of the MAPK pathway by week 4, whereas activation of cell differentiation-related with signalling pathway Wnt/β-catenin and/or BMP signalling was associated to animals in caged conditions especially for Non-Piezo and injury groups (Fig. 7c). Critically, tendons repaired with piezoelectric scaffolds and/or subjected to mechanical loading demonstrated significant upregulation of TNMD or SCX together with significant activation of MAPK/ERK pathway (pERK/tERK). Following 2 weeks post-injury, KCNK2 (Fig. 7c,d, Fold-change ± IQR; Injury=2.03±0.17, p<0.01), TRPA1 (Injury=1.63±0.34, p<0.05) and PIEZO2 (Injury=1.81±0.18, p<0.05) were significantly modulated and were associated with SCX (Injury=2.45±0.25, p<0.01) and TNMD (Injury=1.95±0.37, p<0.05) upregulation.

Protein array analysis (Fig. 7c,d; Fold-change ± IQR) indicated that the fold-change of collagen II synthesis, (Injury=2.04±1.8, p<0.05; Non-piezo=3.20±3.2, p<0.05 and; Piezo2.=3.6±2.08, p<0.01 at week four and Injury=4.01±4.01, p<0.05 at week eight) and BMPR expression (Non-piezo=2.35±1.2, p<0.001 at week four with respect to caged controls) between Tr and Cage groups relative to two weeks were significantly upregulated and associated with significant activation of the BMP signalling pathway via phosphorylation of Smad 1/5/8 at week eight (Injury=2.34±0.3, p<0.05 and; Non-piezo=2.72±2.72, p<0.05 with respect to caged controls). Conversely, upregulation in the expression of MAPK signalling pathway via phosphorylation of ERK1/2 (Piezo=2.23±0.2, p<0.001 at week two with respect to caged control) together with significant regulation of BMP signalling for EMS (Piezo=1.33±0.7 with respect to Injury=4.48±4.48, p<0.05 at week two) was obtained. Notably, after eight weeks, significant activation of BMP signalling was observed only for mechanical stimulation groups (Injury=2.34±0.3, p<0.05 and; Non-Piezo=2.73±2.73, p<0.05 with respect to caged controls). In addition, mechanical stimulation resulted in activation of BMP signaling (Injury=2.35±2.35 with respect to cage control, p<0.05), BMPR upregulation (Non Piezo=2.34±1.39 with respect to cage control, p<0.01) and increased collagen II synthesis (Non-Piezo=3.20±3.20 with respect to cage control, p<0.01) at week four with respect to caged controls. The sustained activation of BMP signaling starting at week two (Injury=4.48±4.48, p<0.05 and Non-Piezo=2.72±2.72,p<0.05 with respect to caged controls) was associated with significant immediate activations of TRPA1 (Injury=1.64±0.34 at two weeks with respect to two weeks caged control, p<0.01 and; Non-Piezo=2.96±1.41, p<0.05 at four weeks with respect to caged group and four weeks Injury=1.44±1.30,p<0.01 and four weeks Non-Piezo, p<0.001 with respect to eight weeks Injury=0.47±0.06),TRPV1 (Non-Piezo=1.91±1.91 at four weeks with respect to Non-Piezo=0.72±0.25 at 2 weeks, p<0.01 and Non-Piezo=0.75±0.65 at eight weeks, p<0.01), KCNK2(Injury=2.02±0.17, p<0.01 at week 2 with respect to caged group) and Piezo2 (Injury=1.81±0.18, p<0.01 at two weeks and four weeks Non-Piezo=1.81±1.81, p<0.05 with respect to eight weeks Injury=0.41±0.41). Conversely, early activation of MAPK (Piezo =2.23±0.40, p<0.01 at week two with respect to caged group) together with decreased levels of BMPR (Piezo=0.55±0.19 at week two with respect to control group and; Piezo=0.77±0.18 at week four with respect to Non-Piezo=2.35±1.51, p<0.01), reduced collagen synthesis II (Piezo=0.66±0.38 at week two and compared to Non-Piezo=3.20±3.20 at week four, p<0.01) and reduction of BMP signaling (Piezo=1.33±0.70 with respect to Injury=4.48±4.48 at week two, p<0.05) was associated with decreased levels of ion channel expression including TRPA1 (Piezo=1.32±1.10 with respect to Non-Piezo=2.96±1.41 at week four, p<0.01), Piezo2 (Piezo2=0.35±0.22 with respect to Non-Piezo= 1.81±1.81 at week four, p<0.01), TRPV1 (Piezo=0.68±0.13 at week four with respect to Non-Piezo=1.91±1.91 at week four, p<0.01) and KCNK2 (Piezo=0.69±0.26 at week four with respect to Piezo at week two, p<0.01).

Evidence suggests that the balance between Ca^2+^, BMP, Wnt/β-catenin and MAPK under biophysical stimulation (mechanical or electromechanical) can control cell dedifferentiation and functional tissue repair processes ^92,93^ (Fig 7d). It is well established that K2P channels are regulators of pain signaling. Both KCNK2 and KCNK4-deficient mice (including double knockouts) show hypersensitivity to mechanical stimuli and under inflammatory conditions (colitis), a downregulation in the expression of KCNK channels results in cellular hyperexcitability^94^. These studies demonstrate that KNCK ion channels can mediate biological functions by regulating intracellular calcium activity. Studies have also shown that different electrical stimulation conditions can selectively influence cell activity and determine cell fates processes. For instance, high levels of electrical stimulation (45Hz) have been observed to increase osteogenesis of MSC through increased expression of RUNX2, ALP and Ca^2+^. In contrast, low-level electrical stimulation (7Hz) promoted osteoclast MSC differentiation via MAPK signalling pathway activation^95^. This study demonstrates that electrical stimulation increases cytoplasm calcium activity and accelerates differentiation processes via regulation of expression of ion channels^96,97^.

### Outlook

This work shows that functional tendon tissue repair and remodelling can be modulated by mechanical and electromechanical stresses. Electromechanical stimulation was enabled by the use of a piezoelectric scaffold and represents a paradigm shift in the management of severe tendon injuries or calcific tendinopathy without the use of drugs or external stimulation. Importantly, this study establishes the engineering foundations for a broad range of synthetic piezoelectric scaffolds that enable bioelectronic control of specific tissue regenerative processes.

## Materials and methods

### Cell culture

Human tendon cells, which were isolated from two donors 63 and 66 years old (LOT #TENM012214F and #TEN020415L), were purchased from Zen-Bio (Research Triangle Park, NC). Cells were expressing TBSH4 and surfaces markers were characterized (CD90+/CD29+; CD45-/CD31-). Additional Human tendon derived cells were harvested from patellar tendon during tendon grafting operations after obtaining written informed consent in the University Hospital Galway From these tissue specimens, primary human tenocytes were isolated and cultured. They were identified as tenocytes through their characteristic growth pattern and by detection of scleraxis (SCX) and tenomodulin (TNMD) expression. TDCs were cultured in Dulbecco’s Modified Eagle’s Medium (DMEM/F-12 with Glutamax, Gibco-BRL) supplemented with penicillin (100 U ml-1), streptomycin (10μg ml-1) (both Sigma-Aldrich) and 10% fetal calf serum (Gibco-BRL). TDCs were cultured in an incubator set at 37°C with 5% CO2 environment and subcultured to passage 2-3 before use. At least 3 donors were used for all assays. Culture media was changed every 3 days.

### Electrospinning

P(VDF-TrFE) and PTFE were made into fine fibres by varying the voltage, mandrel rotation speed and the distance between tip and collector with a constant flow rate. Each polymeric solution was placed in an Aladdin programmable syringe pump (AL-1000) (World Precision Instruments) and attached to a 27-gauge stainless steel needle. An aluminium foil sheet was cut into 10 cm x 28 cm rectangles and taped around the cylindrical mandrel collector (Linari Engineering S.r.l). P(VDF-TrFE) 75/25 weight % (Solvay Solexis) was dissolved in a 3:2 volume ratio of dimethylformamide (DMF)/acetone (Sigma-Aldrich) at a polymer-solvent concentration of 15-35% w/w. Briefly, 2 ml of solution was placed into a 10 ml plastic syringe (Luer lock). Any excess air remaining in the syringe was removed, and a stopper was placed on the syringe to prevent the sample from drying out. The syringe was placed 6 cm away from the collector, the flow rate was constant at 1 ml/hour, and the collector speed was set at 3700 rpm. Finally, the voltage was switched on and increased up to 24-26 kV. The collector was also charged with −6 kV. The experiment ran for 2 hours, and lower humidity (40-50%) yielded more favourable results. PTFE was electrospun with a 2 step process. First, 240 milligrams of poly(ethylene oxide) (PEO), average Mv □300,000 (Sigma-Aldrich), was weighed out and mixed with 10 ml of PTFE, 60 wt % dispersion in H2O (Sigma-Aldrich). The syringe was clamped into a polymer blender and mixed gently for 2 hours. After blending, the air from the syringe was removed using a centrifugation tube at 1500 RPM for 2 minutes. The solution was removed from the centrifuge and filtered through a syringe filter with a 25 mm diameter (Merck Millipore) into a clean syringe at least twice to ensure no precipitates were present. The syringe was placed 10-12 cm away from the collector, and the flow rate was pre-set at 1.25 ml/hour. The rotator was set at the maximum velocity of 2000 RPM since fibres at higher speed tend to break due to their poor mechanical integrity. The voltage was set to 11/12 kV. The experiment ran for 4-5 hours, and a higher relative humidity (50-60 %) yielded more favourable results. Finally, PTFE mats were collected and annealed at 185°C for 5 minutes.

### Fibronectin functionalisation of electrospun scaffolds

Post electrospinning, due to sample chemistry the sample material was functionalized before cell seeding. The samples were carefully removed from the aluminium foil and placed on a square sample holder. Three functional chemistries were investigated to aid in the promotion of cell attachment: Collagen (50 μg/ml), Poly-L-lysine (PLL) (10 μg/ml) and fibronectin (FN) (20 μg/ml). The accessibility of the cell-binding domain was ensured by covalent binding using Poly-Acrylic Acid (PAAc) as a spacer.

### X-ray diffraction (XRD)

X-ray diffraction patterns from films were collected by using a Philips Xpert PRO automatic diffractometer operating at 40 kV and 40 mA, in a theta-theta configuration, secondary monochromator with Cu-K-alpha radiation (λ = 1.5418 Å) and a PIXcel solid-state detector (active length in 2θ 3.347°). Data were collected from 5 to 70° 2θ (step size 0.026° and time per step = 347 s) at RT. A fixed divergence and anti-scattering slit giving a constant volume of sample illumination were used. The signal deconvolution was evaluated using the peak-fit option of the WinPLOTR program. The profile fitting procedure (XRFIT calculation) uses pseudo-Voigt functions with a global FWHM (Full width at half maximum and a global eta (proportion of Lorentzian), and a linear background. Each peak is characterised by its position, intensity, FWHM and eta shifts concerning the global parents. The average size of the crystalline domains β (coherently diffracting domains) of the samples was extracted from the broadening of the signal using the Scherrer equation:

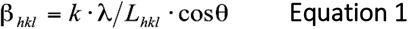

Where βhkl is the broadening of the diffraction line measured at half the maximum line intensity (FWHM) taking into account instrumental contribution (βlnst=0.1°), λ is the X-ray wavelength, Lhkl is the crystal size, and θ is the diffraction angle. K is the Scherrer shape factor (k=0.9 was used for the calculations).

### Differential scanning calorimetry (DSC)

The thermal transitions of the PVDF-TrFE scaffolds were determined on a DSC Q200 (TA Instruments). Samples of 8-10 mg were heated from −10°C to 200°C at 10°C min-1. This first scan was used to determine the melting temperature (Tm) and the melting enthalpy (ΔHm), as well as the Curie temperature (Tc) and the Curie enthalpy (ΔHc). After this first scan, the samples were quenched in the DSC and a second scan was collected from −10 to 200°C at 10°C min-1. For non-isothermal crystallisation studies, the films were heated at a constant heating rate (30°C /min) from room temperature to 260°C and held there for 2 min to eliminate the residual crystals and memory effects due to thermal history, and then cooled to room temperature to crystallize at the same cooling rate under a nitrogen environment. For isothermal crystallisation kinetics, the films were heated at a constant heating rate of 30°C/min from room temperature to 260°C and held there for 2 min to eliminate the thermal history, then the melt was cooled at the same rate up to 148°C and kept constant at 148°C for 10 min until the sample completely crystallized.

### Fourier transform infrared (FTIR)

Infrared spectra of PVDF-TrFE scaffolds were recorded on a Nicolet AVATAR370 Fourier transform infrared spectrophotometer (FTIR) operating in the Attenuated Total Reflectance (ATR-FTIR) mode. Spectra were taken with a resolution of 2 cm^-1^ and were averaged over 64 scans. Using the absorption band of α and β phases at 532 cm^-1^ and 846 cm^-1^, the fraction of β phases were calculated using the following equation The β-phase content F(β) was calculated by the following equation 1:

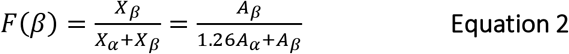

where AEA is the absorbance value at 841 cm^-1^, Aα is the absorbance at 764 cm^-1^, and the factor 1.26 is the ratio of absorption coefficients at 841 cm-1 (K841 = 7.7 x 104 cm2 mol^-1^) to 764 cm^-1^ (K764= 6.1 x 104 cm2 mol^-1^) at the respective wavenumber.

### Mechanical properties

Tensile analysis of scaffolds was performed using a Zwick/Roell Z010 with a 1 kN load cell with a crosshead speed of 500 mm/min and a maximum extension of 500%. Thin scaffolds were cut per ASTM D 882 type specimen. Results are presented as average values of n = 5 replicate experiments with standard deviation.

### Fibre alignment

The samples were mounted onto SEM stubs using double sided conductive tape They were then coated with Gold/ Palladium (approximately 10-20 nm) using a SEM coating system (, UK). The samples were viewed on a SEM running at 5 kV. Fibre alignment was analysed by OrientationJ plugin from ImageJ (free download from NIH).

### Voltage outputs

To analyse the electromechanical behaviour of the scaffolds, a custom made rig was developed to monitor applied strain and induced voltage using an electrometer (Keithley 6514) with virtual infinite input impedance. The Keithley electrometer was connected to the scaffold with a triaxial cable possessing three inputs (high, low and ground). However, leakage resistance and capacitance can appear between the inputs and affect the measurement precision by introducing an offset ramp. In this case, a plausible solution was to use guarding to eliminate these effects. In this mode, the electrometer drives a current through the conductive sheet surrounding the sample inducing the same potential as the high input terminal. With both ends at the same potential, current leakage is not possible. The electrometer used here can operate in guarded mode or un-guarded mode, as appropriate. Finally, we used LabVIEW from National Instruments to collect the data and a mechanical loading reactor to apply strain to the samples following a sinusoidal signal at different frequencies and fixed strain.

### d33 measurements PFM

The piezoelectric response of the PVDF thin films was investigated using piezoresponse force microscopy (PFM) (Bruker Nano Inc, Santa Barbara, CA, USA). For PFM imaging, a Pt-Ir coated conductive probe (SCM-PIT) with a spring constant of 2.8 N/m and a resonant frequency of 75 kHz was used. The amplitude of the detected piezoelectric signal is related to the piezoelectric coefficient (d33) of the material, whereas the phase of the signal reflects the direction of the polarisation of the domains. An average of ten different locations on one sample was used to compute the average value of piezo-response amplitude. Before imaging the PVDF film samples, the PPLN (Periodically Poled Lithium Niobate) test sample was used for standard PFM imaging verification. The vertical piezoresponse was calibrated using deflection sensitivity of the AFM cantilever tip obtained from the force-displacement curve.

The d33 measurement procedure is described below:

> d33 (nm/V) = Amplitude (in V) * deflection sensitivity (nm/V) / (vertical deflection gain (16) * Applied AC Bias)

where deflection sensitivity: 114 nm/V, Vertical deflection gain: 16X and applied AC bias: 0.3 V.

### Mechanical Loading Bioreactor

Briefly, cells were cultured on control and experimental scaffolds at a density of 7 500 cells/cm2 and left for 24 hours to attach. Axial mechanical loading was provided via a UniMechanoCulture T6, Cellscale^®^ bioreactor. Cells were kept under static or dynamic conditions for 8 hours/day of 4% deformation (2 mm) at 0.5, 1 and 2 Hz for 24h, five days or ten days. A total of six scaffolds were used for each experiment.

### Immunostaining

After scaffolds were stimulated with mechanical loading at 4% starin, the samples were he samples were fixed in 4% formadehyde 15 minutes at RT. s. After that, permeabilisation buffer was added and incubated at 4°C for 5 minutes. Samples were then blocked with 1% BSA in 1xPBS at 37°C for 5 minutes. Primary antibody TNMD (1:20, ab203676, Abcam) and rhodamine-phalloidin (Invitrogen, Thermo fisher, 1:500) diluted in 1% BSA in PBS were added and incubated at 37C for 2 hour. Finally, samples were washed with PBS/0.5% Tween-20 for 5 min. Secondary antibody (Alexa Fluor 488, Invitrogen) diluted in 1% BSA in PBS was added and incubated at 37°C for 1 hour. Samples were washed again with 1xPBS/0.5% Tween for 3. times (5 minutes each). DAPI ((4’,6-diamidino-2 phenylindole, Invitrogen) was added and samples were covered with coverslips. Images were processed using ImageJ (Version 1.48, USA).

### RT2-qPCR array processing

RNA was extracted using a trizol and chloroform precipitation method and purified using a Qiagen column. RNA quality after purification was assessed using an Agilent 2100 Bioanalyzer and RNA 6000 Pico Kit for low sample quantities (Agilent Technologies, USA) providing data on the integrity of RNA (RNA Integrity Number; RIN) and RNA concentration. RNA concentrations above 20 ng/μl and RIN >9 were used for subsequent conversion into cDNA. Following the manufacturer’s instructions, cDNA was synthesised by reverse transcription of RNA using the RT2 First Strand Kit (SA Biosciences). PCR was performed in iQ5 Thermal Cycler (Bio-Rad, Munich, Germany), using diluted cDNA as template (10 μl of cDNA, 10 μl of Genomic DNA Elimination Mixture and 91 μl of RNase free water) and the RT2 SYBR Green qPCR Master Mix (SA Biosciences), according to the manufacturer’s guidelines. Each experiment was done in triplicate using the two donors (N=2), and all samples were analysed in technical duplicate for each gene set. Quantification results were normalised against stable housekeeping genes (TOP1, GAPDH, HSP90AB1, RPLP0 and TBP), using the geometric mean of the threshold cycle (Ct) of the genes included in each plate. Expression changes were calculated using the ΔΔCt method and followed by calculation of regulation fold changes.

**Table.**
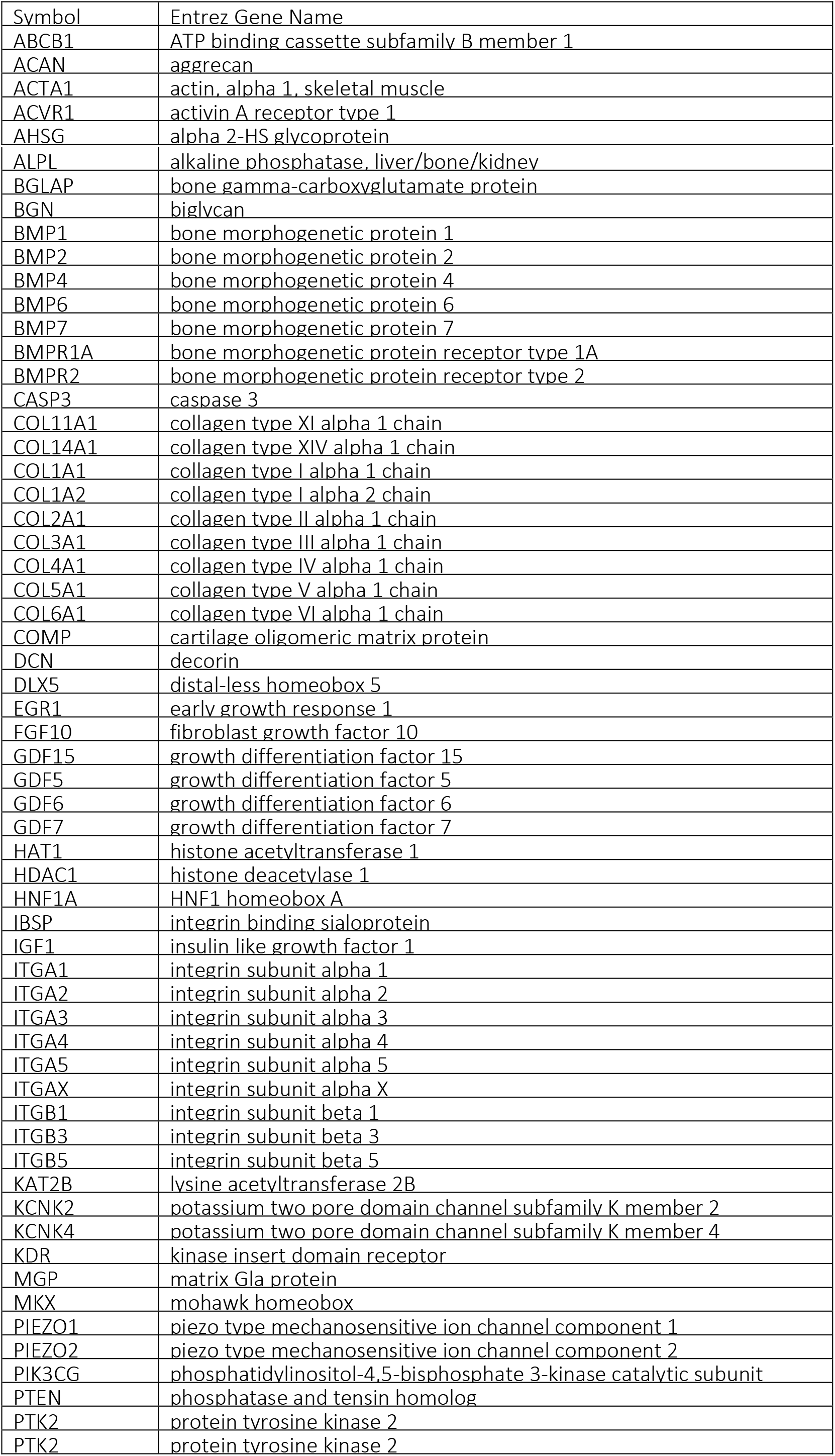
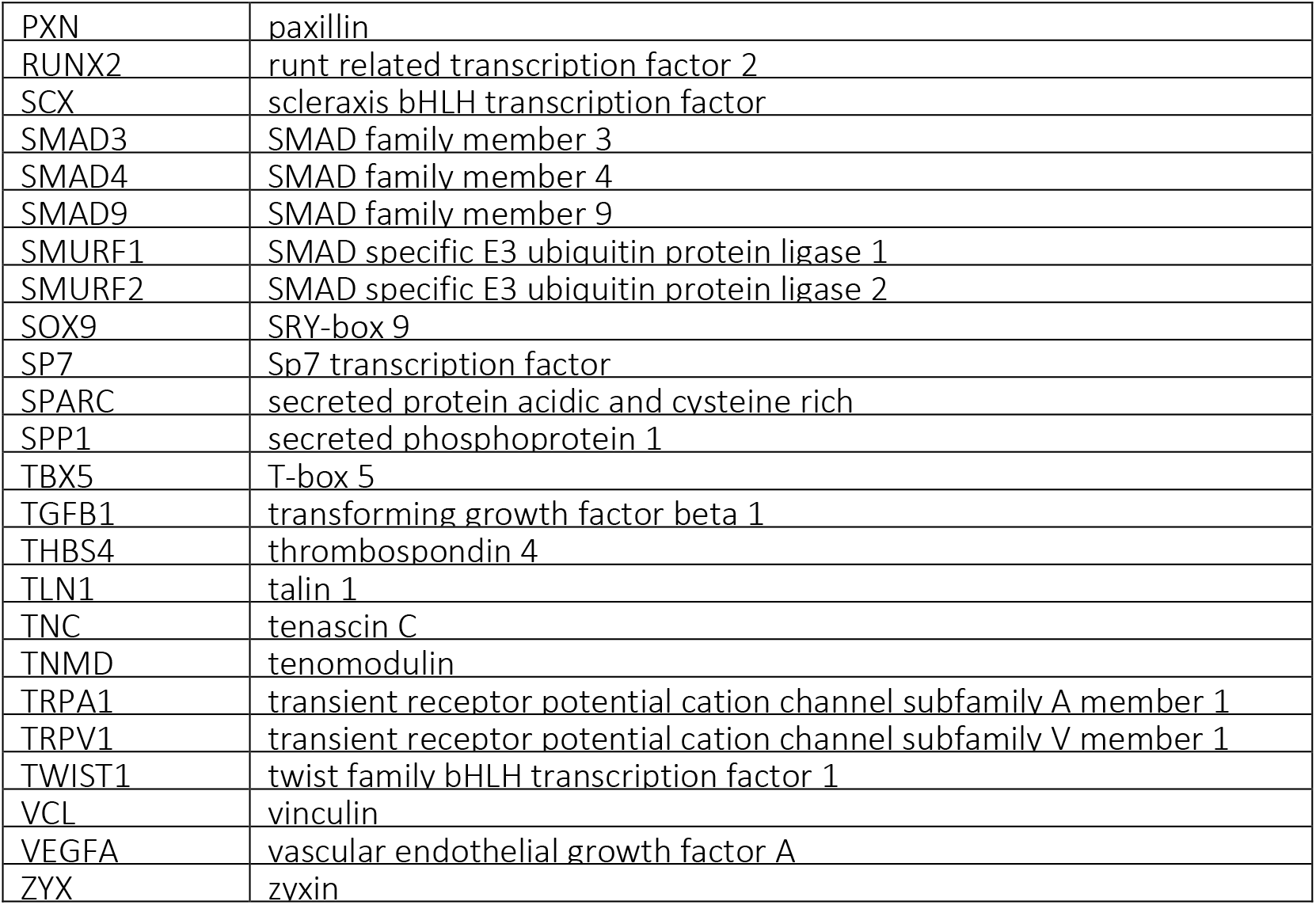
QIAGEN genes list

### Protein Expression

Protein Antibody Microarray was custom made. Nexterion slide H microarray slides were purchased from Schott AG (Mainz, Germany). Alexa Fluor 555 carboxylic acid succinimidyl ester was obtained from Life Technologies (Carlsbad, CA, USA). Protein samples were labeled with Alexa Fluor 555 carboxylic acid succinimidyl ester according to manufacturer’s instructions. The excess label was removed, and the buffer was exchanged with PBS, pH 7.4, by centrifugation through 3 kDa molecular weight cutoff filters. Absorbance at 555 and 280 nm was measured for labelled samples and calculations were performed according to manufacturer’s instructions using an arbitrary extinction coefficient of 100,000 and molecular mass of 100,000 to enable quantification of relative protein concentration and label substitution efficiency. All commercial antibodies (Table 1) were buffer exchanged into PBS and quantified by a bicinchoninic acid (BCA) assay. Antibodies were diluted to print concentration in PBS and printed in six replicates on Nexterion H amine-reactive, hydrogel-coated glass slides using a SciFLEXARRAYER S3 piezoelectric printer (Scienion, Berlin, Germany) under constant humidity (62% +/− 2%) at 20 °C. Each feature was printed using ≈1 nL of diluted antibody using an uncoated 90 μm glass nozzle with eight replicated subarrays per microarray slide. After printing, slides were incubated in a humidity chamber overnight at room temperature to facilitate complete conjugation. The slides were then blocked in 100 × 10-3 m ethanolamine in 50 × 10-3 m sodium borate, pH 8.0, for 1 h at room temperature. Slides were washed in PBS with 0.05% Tween 20 (PBS-T) three times for 2 min each wash followed by one wash in PBS, dried by centrifugation (470 × g, 5 min), and then stored with a desiccant at 4 °C until use. Incubations were carried out in the dark. Microarray slides were incubated as previously described. Initially, one labelled sample was titrated (2.5–15 μg mL–1) for optimal signal to noise ratio and all samples were subsequently incubated for 1 h at 23 °C at 9 μg mL-1 in Tris-buffered saline (TBS; 20 × 10-3 m Tris-HCl, 100 × 10-3 m NaCl, 1 × 10-3 m CaCl2, 1 × 10-3 m MgCl2, pH 7.2) with 0.05% Tween 20 (TBS-T). All microarray experiments were carried out using three replicate slides. Alexa Fluor 555 labelled cells lysate (10 μg mL-1) were incubated in two separate subarrays on every slide to confirm retained antibody performance and printing, respectively (figure 1). After incubation, slides were washed three times in TBS-T for 2 min per wash, once in TBS and then centrifuged dry as above. Dried slides were scanned immediately on an Agilent G2505 microarray scanner using the Cy3 channel (532 nm excitation, 90% photomultiplier tubes (PMT), 5 μm resolution) and intensity data were saved as a.tif file. Antibody microarrays were verified to remain active for at least 2 weeks after printing and all incubations were carried out within that timeframe. Data extraction from.tif files was performed mainly as previously described. Data were normalized to the mean of three replicate microarray slides (subarray-by-subarray using subarray total intensity, n = 4, 24 data points). Unsupervised hierarchical clustering of normalised data was performed using Hierarchical Clustering Explorer v3.0 (http://www.cs.umd.edu/hcil/hce/hce3.html) using the parameters no prefiltering, complete linkage, and Euclidean distance. All data presented here were confirmed using at least four replicates for each of the test groups and control group. The results are expressed as the mean of the values ± standard error of the mean.

Different types of arrays were fabricated to investigated tendon regeneration or phenotype maintenance (tenogenesis), intracellular molecular pathways (signalling) or membrane proteins (receptors).

**Table 1.** *<u>Tenogenesis array</u>*. Proteins associated with tendon regeneration or tenocyte function.

| Proteins | Abbreviation | Purchased from | Cat. no |
| --- | --- | --- | --- |
| Scleraxis | SCX | Abcam | ab58655 |
| Tenomodulin | TNMD | Abcam | ab203676 |
| Byglican | BGN | Abcam | ab49701 |
| Decorin | DCN | Abcam | ab175404 |
| Thrombospondin 4 | THBS-4 | Abcam | ab176116 |
| Tenascin C | TNC | Abcam | ab88280 |
| Collagen I | COLI | Abcam | ab138492 |
| Collagen II | COLI | Abcam | ab185430 |
| Collagen III | COLIII | Abcam | ab7778 |
| Collagen V | COLV | Abcam | ab7046 |

**Table 2.** *<u>Signalling array</u>*. Proteins associate to different signalling pathways (MAPK, FAK, TGF-B, BMP and WNT).

| Proteins | Abbreviation | Purchased from | Cat. no |
| --- | --- | --- | --- |
| Smad 1 | SMAD1 | Cell signalling® | 6944S |
| Smad 5 | SMAD3 | Cell signalling® | 12534S |
| Phosphorylated Smad 1 | SMAD1/5/8 | Cell signalling® | 5753S |
| Phosphorylated Smad 1/5/8 | pSMAD158 | Cell signalling® | 9516s |
| Focal adhesion kinase | FAK | MBL® | D061-3 |
| Phosphorylated FAK | pFAK | Cell signalling® | 3284S |
| MAPK | ERK | Cell signalling® | 4696S |
| p44/42 MAPK | pERK | Cell signalling® | 4377S |
| Wnt/ β-catenin | β-Catenin | Millipore® | 2858901 |
| Active β-catenin, clone 8E7 | Active β-Catenin | Millipore® | ABC |
| β-actin | β-actin | WAKO® | 019-19741 |

**Table 3.** Receptors array. Proteins associate to cell membrane receptors.

| Proteins | Abbreviation | Purchased from | Cat. no |
| --- | --- | --- | --- |
| TRPV1 | TRPV1 | SantaCruz® | sc-20813 |
| Piezo1 | Piezo1 | SantaCruz® | sc-164319 |
| Piezo2 | Piezo2 | SantaCruz® | sc-84763 |
| TRPA1 | TRPA1 | SantaCruz® | sc-32353 |
| KCNK2 | KCNK2 | SantaCruz® | sc-11557 |
| KCNK4 | KCNK4 | Abcam® | ab81367 |
| L-type Ca<sup>2+</sup> | L-type Ca<sup>2+</sup> | SantaCruz® | sc-25686 |
| BMPR1A | BMPR1A | ThermoFisher® | PA5-11856 |
| Integrin1 | ITG1 | Abcam® | ab134179 |
| Integrin3 | ITG3 | Abcam® | ab34409 |
| Integrin5 | ITG5 | Cell signalling® | 3629S |

### In vivo animal model

The animal care research ethics committee at the National University of Ireland, Galway and Health Products Regulatory Authority (AE19125/P055) approved all the animal procedures used in this study. Also, animal care and management followed the Standard Operating Procedures of the Animal Facility of the National University of Ireland, Galway. Animals were allowed to acclimatise for at least seven days before any surgical procedures. Subsequently, animals were acclimated to the treadmill running for one week, and their behaviour was analysed. A total of 105 Female Lewis rats aged 6-8 (220g) weeks were used in this study. The animals were anaesthetised by isofluorane inhalation (5% induction reducing to 1-2% for maintenance during procedures). The right leg was shaved and swabbed with iodine to minimise the risk of bacterial contamination. An incision was created through the skin (~1cm) from the myotendinous junction distally to the osteotendinous junction. The incision provided ample exposure to the Achilles tendon. The fascia surrounding the Achilles was transected longitudinally and carefully retracted to expose th the Achilles tendon. Before implantation and tendon transection, two looped sutured were inserted at the top (muscle) and bottom (bone). After the total tendon length was measured a 3 mm defect in proximal/distal extension (at 3 mm from the calcaneus) was created using a positioning device and an 11 surgical blade, resulting in a 6 mm gap after tissue retraction. The construct was then sutured (4-0 Ethicon) to both ends of the tendon to bridge the gap using a modified Kessler technique, and the skin was sutured. After a period of two, four and eight weeks the animals were euthanised and tendon tissue, as well as contralateral tendons, were harvested. Animals recovered for 2 weeks and were gradually exposed to treadmill running (see Table). Briefly, the treadmill running was increased from once a week for 5 min to 30-45 mins for 5 days a week.

**Table 4.** Exercise protocol used for the rat running group (treadmill)

| Time |  | Duration (min) | Speed (m/min) |
| --- | --- | --- | --- |
| Week 2 | Day 14 | 5 | 8-9 |
|  | Day 15 | 10 | 9-10 |
|  | Day 16 | 15 | 9-10 |
|  | Day 17 | 20 | 9-10 |
|  | Day 18 | 30 | 9-10 |
| Week 3 | Day 19 | 45 | 9-11 |
|  | Day 20 | 30 | 9-11 |
|  | Day 21 | 45 | 9-12 |
|  | Day 22 | 30 | 10-14 |
|  | Day 23 | 45 | 10-14 |
| Week 4 |  | 30 | 10-14 |
| Week 5 |  | 45 | 10-14 |
| Week 6 |  | 30 | 10-14 |
| Week 7 |  | 45 | 10-14 |
| Week 8 |  | 30 | 10-14 |
| Week 9 |  | 45 | 10-14 |
| Week 10-16 |  | 30 | 10-14 |

### Histology

Repaired tendon tissues and contralateral tendons of all groups (n=7) were dissected from the proximal myotendinous junction to the distal osteotendinous junction and processed for histological analysis. The samples were fixed in 10% neutral buffered formalin (24 hours), dehydrated in gradient alcohols, cleared, and embedded in paraffin blocks, as reported previously. Histological sections (6 μm thick) were prepared using microtome sectioning (Leica Rotary Microtome). In order to distinguish between scar tissue or new tendon formation, polarisation microscopy and picrosirius red staining were used. Also, for descriptive histology 6 μm, thick sections were stained using Hematoxylin & Eosin stain, Masson-Goldner’s stain, Alcian Blue, O-safranin stain, Red picrosirius satin or Herovici’s polychrome stain according to the manufacturer’s guidelines. Areas of chondrification within the defect region at 4 or 8 weeks after surgery were measured using Safranin O staining and ImageJ (v1.52i). For each tissue, 3 different frontal-longitudinal sections were analysed (1 section of the middle tendon, 1 section ventral to this middle part and 1 section dorsal to this middle part). For each section 5 consecutive images were captured spanning the entire length of the tissue, omitting the transition zones of original tendon stumps to regenerated tissue. The volume fraction (VV) of tendon cells were used to estimate cell proliferation. A 192-point grid was overlaid on 40X images of H&E stained tissue sections. The number of tendon cells intersecting points of the grid was counted (PP), along with the total number of points on the tissue (PT). The volume fraction of tendon cells (VV) was calculated using the formula below:

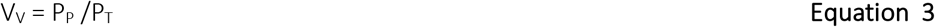

### Histomorphometry and point-based scoring system (composite score)

In order to evaluate the effect of treadmill running on the progression of tendon repair, histomorphometric measurements on stained samples from three animals (5 sections in total per animal) were employed. A point-based scoring system was used, and the following parameters were scored based on the percentage of 1) increase in the number of cells compared to control intact tendon (sham) 2) Increase of calcification compared to intact tendon 3) Increase in the vascularization and innervation compared to intact tendon 4) Increase of fat deposits compared to intact tendon 4) Increase of fiber orientation compared to intact tendon 5) Increase of cell morphology compared to intact tendon.

### Micro-CT scan

At 8 weeks after Achilles tenotomy, the right hind legs of each group were collected at sacrifice for micro-computed tomography analyses (μCT 100, Scanco Medical, Bruttisellen, Zurich, Switzerland). Samples were scanned using the following settings: 60 kV, 150 μA with a mean 20 μm slice thickness. The reconstructed scaffold and Achilles tendons were selected for quantification of calcifications volume.

### Functional recovery analysis

An animal treadmill (Exer 3/6, Columbus Instruments) track integrated with a video-based system was used to obtain spatiotemporal parameters of gait. The animal gait was analysed through a clear plastic Lexan at the sides of the system consisting of a cage (50.8 cm x 50.8 cm x 33 cm) with gates placed at each end of the walkway. A digital camera (8 M pixels and 120 frames per seconds) was positioned 30 cm in front of the walkway to capture the sagittal view of the rat from the walkway. The data were analysed and processed with Kinovea software (v0.8.27). For all groups, the system was calibrated using the same scale bar located in the image and modelled the leg motion using pairs of circle markers. The ankle, knee, hip and MTP joint angles (see Figure 5–4) were measured using the automatic black ink marker feature recognition on the hip, knee, ankle, and 3rd metatarsal head at the four gait stages: initial contact, mid-stance, pre-swing, and mid-swing. The spatial and temporal gait parameters analyzed were step length, angle and cycle time. The walking speed was calculated by dividing the step length by the cycle time. Each angle joint curve was normalized by cycle time before further analysis. To determine the change on gait angle, the maximum difference in amplitude (Ampl.) between initial contact and end of swing was measured during an entire step and was calculated according to following formula:

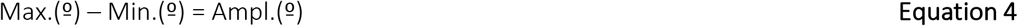

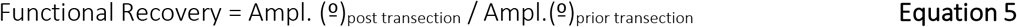

### Statistical Analysis

Statistical analyses were performed using GraphPad Prism software version 8.4.3 (GraphPad Software, CA, USA). To test whether the data was normally distributed normality tests were applied (D’Agostino and Pearson). When data followed a normal distribution a one-way ANOVA analysis for the comparison of means between different groups was performed. Homogeneity of variances was tested using Bartlett’s tests, and in a case of unequal standard deviations (SD), Welch ANOVA test was applied. If data were not normally distributed, the comparison of medians between different groups was assessed by non-parametric Kruskal-Wallis test followed by Dunn post-hoc test. Since most of the data was not normally distributed, most of the results are expressed as median as central tendency characteristic and interquartile range (IQR) or range as dispersion characteristic and, p values of < 0.05 were considered statistically significant.

## Author Contributions

MF-Y, MJPB, AP, MK conceived the experiments. MF-Y, AT performed the experiments. A-T, AS, AL performed research. MK, AP, TS, MP provided materials and expertise. MF-Y, MK analysed the data. AP, MJPB provided critique and context for the data. MF-Y wrote the manuscript. MF-Y prepared the figures. All authors read and commented on the manuscript.

## Acknowledgement

The work was supported by grants to MJPB from Science Foundation Ireland (16/BBSRC/3317). We thank Dr. Oliver Carroll for technical assistance. SGIker technical services (UPV/EHU) are gratefully acknowledged for XRD and XPS support.

## Conflicts of interest

There authors have no conflicts of interest

## Notes

### Competing Interest Statement

The authors have declared no competing interest.

