## Supplemental Information for "Self-powered piezo-bioelectronic device mediates tendon repair through modulation of mechanosensitive ion channels"

### Supplementary information

#### Design of scaffold

There are several advantages of using rotating collector extreme linear speeds during electrospinning. First, it enables the formation of mesoscopic joints between adjacent fibres as a result of residual (not evaporated) solvent that maximise the mechanical properties of the scaffold. Limited evaporation is a function of short flight times and use of high boiling point solvents (i.e. DMAC) as described by Persano et al. in similar studies. And secondly, individual fibres (fibre diameter) resulting in finer and more crystalline fibres and increases the piezoelectrical performance. The use of DMAc in combination with acetone (50:50 v/v) allowed the surface morphology of the fibres to be slightly rough.

As shown in Fig. SI1, oxygen (O1s) and nitrogen (N1s) XPS characteristic peaks confirmed successful fibronectin coating on the scaffold surface. The spectrum of the C1s carbon signal (see Figure 11) at 284.6 V revealed changes in the C-C and C-H bond energy suggesting new covalent bonds have been formed. The other three contributions are related to intra-chain polymer bonds C-F. Finally, the evident Fluor signal present on both spectra suggests that the modification with fibronectin is in the nanoscale range (< 20 nm).

The thermal transition properties of P(VDF-TrFE) scaffolds were evaluated using differential scanning calorimetry (DSC) (table SI1). PVDF-TrFE scaffolds demonstrated two main transitions: an initial endothermic peak, associated with the Curie temperature (Tc ~117ºC) and a second endothermic peak that is associated with the melting temperature (Tm~141ºC). The observed ΔHc and the ΔHm values were ~5 J g-1 and ~17 J g-1 for the scaffolds without annealing. This process modified the thermal properties, and a new endothermic transition occurred at 90ºC (related to pAAC), and the $\Delta H_m$ value decreased to $\Delta H_m$=12.74 J $g^{-1}$. This change is associated with the presence of fibronectin bound to the scaffold.

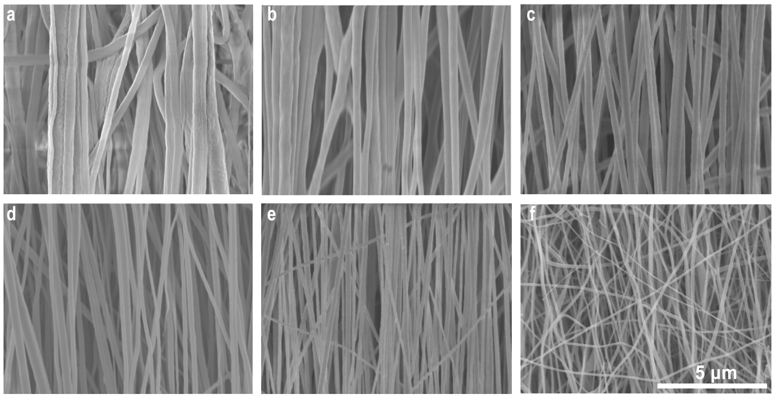


**Supplementary Fig. | 1. a,** Fibers obtained with different polymeric solution concentrations (1.7, 1.4, 1.2 mg/ml respectively) using either long-distance (a,b,c) or short-distance (d,e,f) electrospinning. Fibre diameter and alignment of fibres showed dependency with polymeric solution concentration.

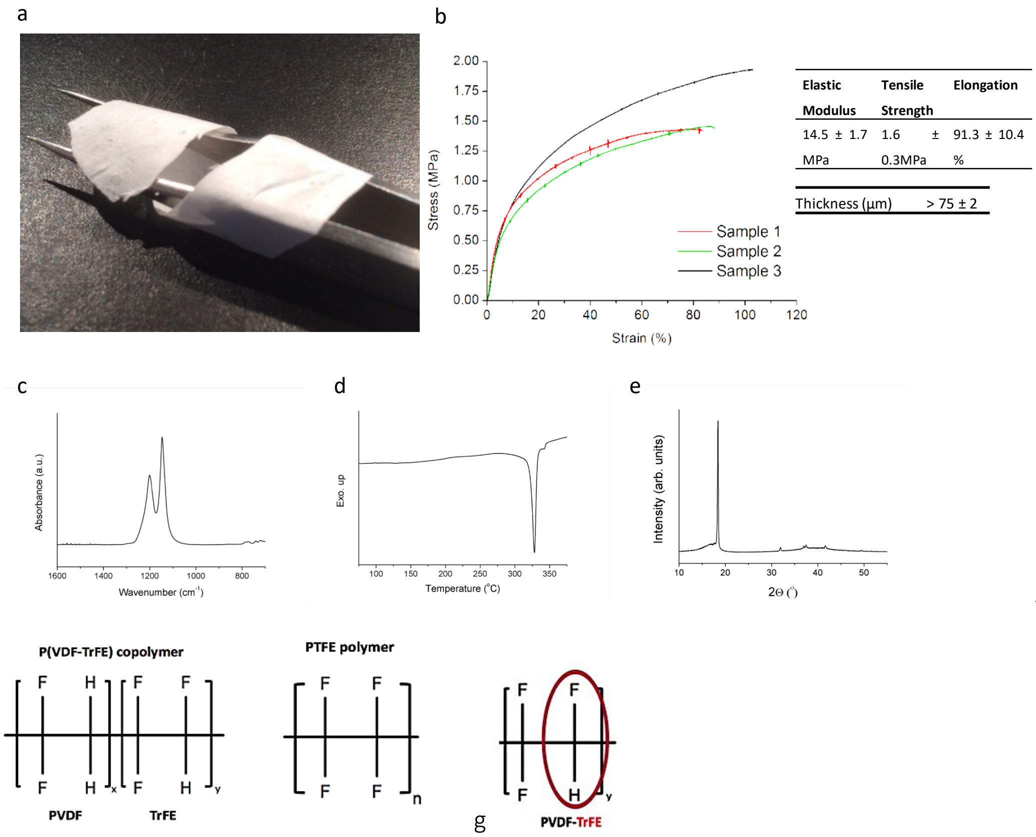

**Supplementary Fig. | 2. a**, PTFE scaffold image. **b**, Mechanical properties of PTFE. **c**, FTIR measurement showing C-F bond. **d**, DSC measurement showing the melting point at 330 C **e**, Schematic of the molecular structure of P(VDF-TrFE) copolymer and PTFE polymer. Note the structure of PTFE with four polar covalent Carbon—Fluorine bonds making it extremely stable and therefore Non-Piezoelectric. **g**, Within the P(VDF-TrFE) copolymer, the -TrFE component exhibits a strong dipole across the Hydrogen to Fluorine bond (red circle).

One major distinction between PE and PTFE chemical structures is the change of C-C chain flexibility. PE has a planar zigzag conformation offering high flexibility to the structure. Conversely, PTFE offers some stereochemical constraint. In crystal form, the chemical conformation of PTFE always results in a helix whereas in PVDF-TrFE due to the ability of chain rotation it sometimes forms helices from the planar zigzag. To attain piezoelectric behaviour, chain rotation for dipole orientation to achieve inherent dipole moment is essential. Also, the partial charges between the Carbon and Fluorine atoms ($C^{\partial+}$—$F^{\partial-}$) attract, generating the 'strongest bond in organic chemistry' (100kcal $mol^{-1}$ bond strength)[98]. For these reasons PTFE is exceptionally chemically stable and exhibits <u>no piezoelectric effect</u>

Direct electrospinning of PTFE is not possible. PTFE is a high-molecular-weight polymer, and the only practical solvent would be a low molecular weight polymer of itself such as a perfluorinated solvent (PFC) containing only C-F bonds and C-H bonds. Due to the strong C-F bond (melting point of 327ºC), dissolving PTFE and obtain a conductive polymeric solution using an organic solvent is not possible. As observed in Fig. SI2c, DSC measurements showed a clear single peak of a highly crystalline phase (ΔXc= 61.1 %, PTFE). FTIR and XRD measurements further confirmed the PEO removal. After sintering, only peaks at 1201 and 1145 $cm^{-1}$ in the FTIR spectra corresponding to C-F bonds from PTFE were obtained. XRD measurements showed four peaks, a unique dominant peak at 18.2 and peaks at 31.8, 37.1 and 42.2 that correspond to the same fingerprint of pure PTFE powder. Mechanical properties obtained showed low mechanical ultimate strength obtained does not correspond to values in the literature it can be attributed to defects presents in the fibres due to PEO decomposition.

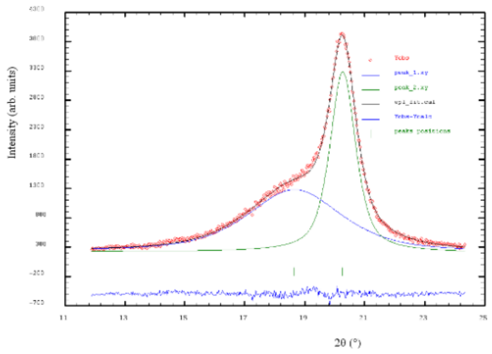


**Supplementary Fig. | 3.** Deconvolution of the XRD spectra into crystalline phase and amorphous halo.

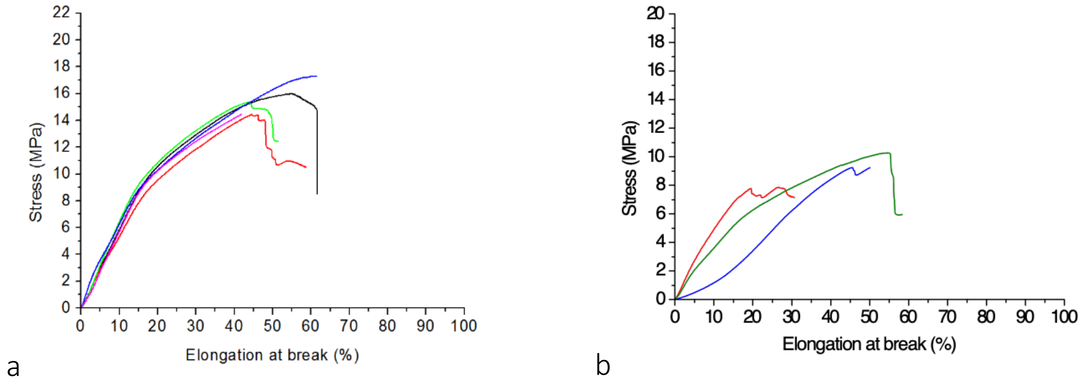


**Supplementary Fig. | 4** Tensile mechanical test spectra for **a**, Short distance (6 cm) **b**, long-distance (20 cm) electrospun scaffolds. Results showed that short distance scaffolds have superior elastic modulus, yield stress, elongation and reproducibility of than long-distance scaffolds.

**Supplementary Table 1.** Degree of crystallinity and the average size of the crystalline domain of the PVDF-TrFE scaffolds.

| *Sample ID* | *Deg.crystallinity %* | *Crystalline size domain* |
|---|---|---|
| Long-distance (random). | *40* | 9 |
| Long-distance (aligned). | *43* | 9 |
| Short-distance (aligned) | *49* | 9 |
| Short-distance (aligned) + annealing | *47* | 11 |

Supplementary Table 2. DSC measurements results. Effect of distance and annealing during surface functionalisation on the thermal properties.

| Sample ID | Diameter | Tc | ΔHc | Tm | ΔHm |
|---|---|---|---|---|---|
| Short-distance | 540 nm | 117 | 5.4 | 140 | 17.1 |
| Long-distance | 540 nm | 117 | 5.2 | 140 | 17.4 |
| Short-distance + annealing | 540nm | 117 | 3.4 | 139 | 12.7 |

Table SI3 shows the results from analysing the FTIR spectra of PVDF-TrFE scaffolds. The bands that are exclusively associated with β-phase were discerned (i.e., bands at 840 cm-1 and 1279 cm-1). However, bands associated with the α-phase (i.e., bands at 764 cm-1 and 976 cm-1) did not appear in the spectra, suggesting that the β-phase is the predominant crystalline phase in the PVDF-TrFE scaffolds. The degree of crystallinity was calculated by deconvolution of XRD spectra into crystalline peaks and amorphous halo. (figure SI3 and table SI3). The samples showed different degree of crystallinity (40-49%) and similar average size of the crystalline domain (~10 nm). Scaffolds spun using long-distance (between collector and needle) showed the lowest β-phase content (58%) compared to short-distance spun scaffolds+annealing (78%).

**Supplementary Table 3.** β-phase content in the function of processing.determined by FTIR.

| Sample ID | Beta phase content / Total |
|---|---|
| Long-distance | 58 |
| Short-distance | 63 |
| Short-distance + annealing | 78 |

In general, thermal annealing (over Tc temperature 132ºC for 2h) resulted in highly crystalline samples (>70% ). The increase in crystallinity after annealing originates from increased γ-phase (15% on annealed samples compared to 1% on as-spun samples). However, under SEM visualisation (data not shown), it was noticed that thermal annealing resulted in significant loss of fibre organisation and introduced defects on the fibre surfaces (in the form of cracks) due to thermal expansion coefficient changes between crystalline and amorphous phases. It was obtained that annealing at 90ºC for 1 hour process showed a significant increase in the crystallinity without affecting fibre structure and surface. Higher annealing temperatures (i.e.110 to 135) had a negative impact on essential characteristics of the samples (fibre organisation and surface morphology, increased brittleness) for the indented tissue engineering application (decreased porosity as fibre form several mesoscale and macroscale joints between the fibres and the total volume of scaffold was reduced by 30%). Interestingly, despite no significant difference in crystallinity level was observed between stretched and the as-spun samples, mechanical properties such as stiffness was significantly improved.

First, a stable dispersion (surfactant content 7.2% wt %) of 160 nm PTFE particles (62 wt%, pH 10.5) was electrospun in combination with a fibre forming agent, polyethene oxide (PEO). Aligned PTFE nanofibres of 690±80 nm diameter were obtained using a ratio between 0.01 PTFE:PEO. Secondly, the scaffolds were annealed at a temperature beyond the melting point (395ºC) producing high-quality and continuous PTFE fibres. Similarly to PVDF-TrFE , prior to cell experiments and for facilitating cell attachment, PTFE surface was chemically treated to covalently bind fibronectin through a nanometric layer of polyacrylic acid.

The diameter of fibres was measured by SEM and showed a reduction from 540 to 516 nm after cold drawing at 12 % strain and correlates well with the theoretical values :

$$V_o = \pi . l_o . d_{o2}/A \qquad \text{Eq. 1}$$

Considering the volume constant and an elongation of 12% the reduction coefficient was 0.95, and the calculated fibre diameter was 513 nm, close to the experimental value.

<u>*In vitro* characterisation</u>

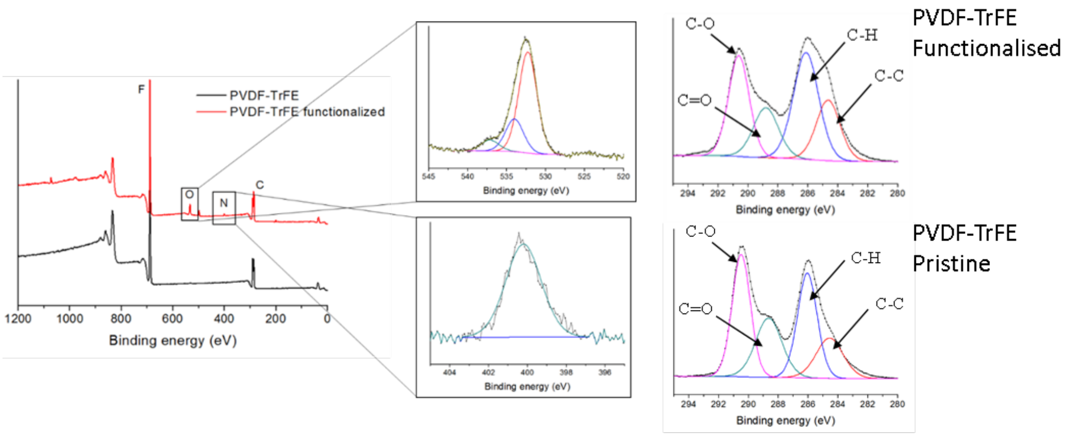


**Supplementary Fig. | 5.** XPS measurements after fibronectin surface functionalisation. Surface chemistry analyses. X-ray Photoelectron Spectroscopy (XPS) spectra are shown for surfaces following fibronectin surface immobilisation (in red) and compared to pristine surfaces without surface functionalisation (in black). High-resolution spectra are showing the presence of peaks corresponding to Nitrogen (N1s at 405.5 eV) and Oxygen (O1s at 530.9 eV). Surface chemistry analyses. X-ray Photoelectron Spectroscopy (XPS) high-resolution spectrum for Carbon (C1s) showing typical hydrocarbon contamination with C==, C-C, C-H and C-O components. Carbon signal was deconvoluted for a) pristine and b) functionalised scaffolds showing significant changes in C-C component.

Under continuous mechanical stimulation of scaffolds, it was observed that cell proliferation was reduced and proportional to the loading frequency. A final stimulation frequency of 0.5 Hz was chosen based on an analysis of cell survival under dynamic conditions (see Fig. 4). It was obtained that tenocytes that lost their elongated morphology underwent significant down-regulation in TNMD and SCX expression when cultured on 2D planar films. Conversely, when dedifferentiated tenocytes were cultured on 3D electrospun scaffolds, they maintained a spindle-like morphology and demonstrated increased expression of TNMD and SCX relative to planar 2D films.

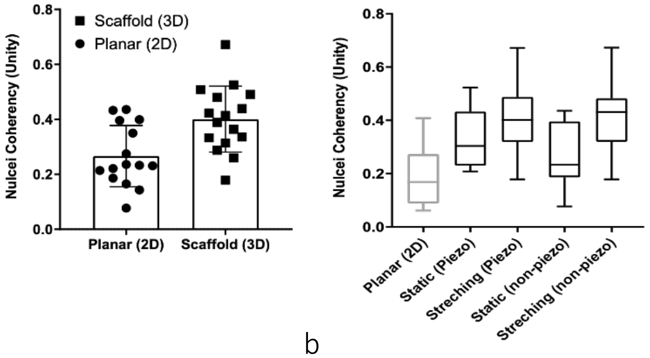


a b

**Supplementary Fig. | 6.** a) Comparison between planar and fibrous (aligned fibres) morphologies on nuclei coherency (aspect ratio/deformation) and b) Comparison between static and dynamic (stretched 4%) cultures of tenocytes using piezoelectric and non-piezoelectric scaffolds on nuclei coherency (aspect ratio/deformation).

A custom-made microarray was designed to assess the expression of genes associated with focal adhesion proteins, collagens, receptors (integrins, ion channels and growth factor receptors) and, inflammation and cell differentiation towards the bone, cartilage or tendon. Gene analysis was performed on cells cultured on non-piezoelectric and piezoelectric scaffolds under mechanical stimulation (0.5 Hz, 8 hours/day) at days one, five and ten.

SCX and MKX were upregulated at days ten and ten and are involved in collagen synthesis and fibrillogenesis. Collagens, predominantly Col1a, constitute the bulk of mature mammalian tendon ECM. Collagen I, III and IV were also upregulated at days five and ten under electromechanical stimulation.

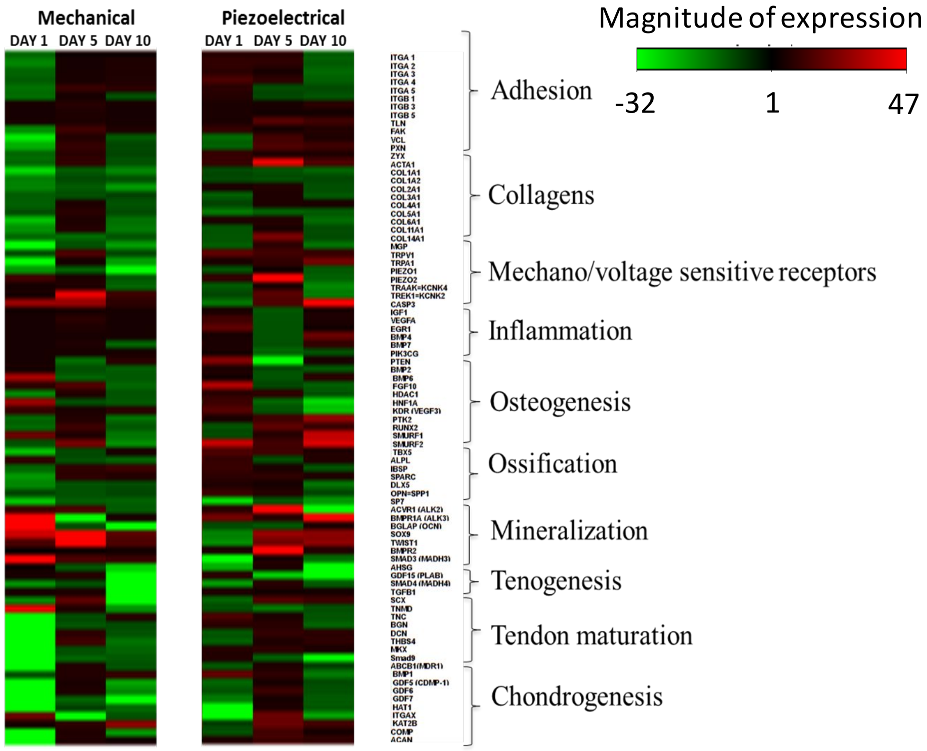


a

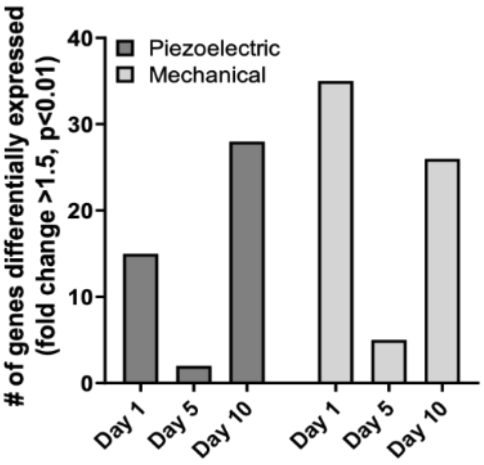


b

**Supplementary Fig. | 7. a**, Gene expression arrays for mechanical and piezoelectric stimulation. **b**, Comparison of genes differentially expressed for mechanical (A) and piezoelectric stimulation (B) at days one, five and ten respectively.

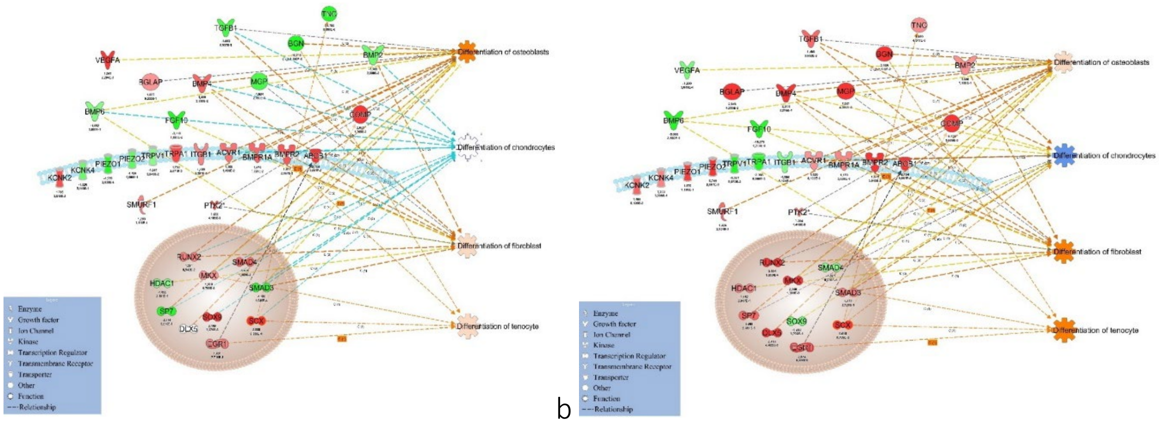


a b

**Fig. | SI8**. Ingenuity pathway analysis. **a**, Mechanical and **b**, electromechanical (piezo) stimulation for ten days induced modulated cell function, including differentiation. Ingenuity pathway analysis identified a mechanistic network of 37 genes which underwent statistically significant modulation. Network analysis of the identified genes: recently discovered mechanoreceptors Piezo1 and Piezo 2 indicated a positive correlation with changes in gene expression and increased tenospecific function.

Tenocytes subjected to mechanical stimulation exhibited down-regulation of the integrin signalling pathway (Fig. 4), with exceptions observed in cytoskeleton rearrangement. Down-regulation of vinculin (VCL) was prominent with mechanical stimulation as was the gene associated with the linker protein paxillin (PXN). Conversely, up-regulation of VCL and PXN was observed in tenocytes subjected to piezoelectric stimulation, initiating cytoskeleton rearrangement mediated transcription of tendon specific proteins. These results were validated using a protein array, and focal adhesion paxillin was highly expressed when comparing EMS and mechanical stimulation (MS). Protein expression was used as a validation method to confirm the differential modulation of osteospecific pathways under EMS and MS. The expression of tendon-related proteins TNMD, TNC and BGN, were upregulated under EM stimulation but downregulated after mechanical stimulation for ten days. The activation osteospecific signalling pathways β-catenin and BMP were investigated through the phosphorylation of SMAD1/5/8 (BMP signalling) and active β-catenin. As shown in Fig. 4, EM stimulation resulted in downregulation of BMP signalling but upregulation of functional proteins including TNMD, TNC and BGN.

*In vivo* characterization

**Supplementary Table 4.** Functional recovery analysis over a period of 8 weeks in animals undergoing treadmill running. Significant differences were observed between the animals undergoing treadmill running relative to animals subjected to static conditions (N=7).

| | | Injury | | | | Non-piezo (static) | | | | Piezo (static) | | | |
|---|---|---|---|---|---|---|---|---|---|---|---|---|---|
| | Ctrl (º) | Static (4 w) | TR (4 w) | Static (8 w) | TR (8 w) | Static (4 w) | TR (4 w) | Static (8 w) | TR (8 w) | Static (4 w) | TR (4 w) | Static (8 w) | TR (8 w) |
| MTP | 100±2 | 52±5 | 48±3 | 51±7 | 71±8 | 56±5 | 58±5 | 55±9 | 92±8 | 55±5 | 48±5 | 52±8 | 62±8 |
| Knee | 80±3 | 53±4 | 58±7 | 63±7 | 67±7 | 55±4 | 62±3 | 55±2 | 72±3 | 56±10 | 60±7 | 53±6 | 65±6 |
| Hip | 18±2 | 12±4 | 15±5 | 9±4 | 15±5 | 10±1 | 14±2 | 12±5 | 15±3 | 10±3 | 8±2 | 8±4 | 12±8 |
| Ankle | 70±2 | 57±11 | 55 ±3 | 58±7 | 61±3 | 53±7 | 54±4 | 51±5 | 67±3 | 55 ±9 | 57±3 | 57±5 | 69±7 |

Figure 5-1. Functional recovery analysis following injury in animals treated with non-piezo and piezo scaffolds (static and TR) at weeks 4 and 8 after injury. It was observed that animals treated with a non-piezo scaffold (TR) showed significant functional recovery in the ankle, MTP and knee joints after 4 and 8 weeks. Results are expressed by mean ± standard deviation, * p<005, ** p<0.01.

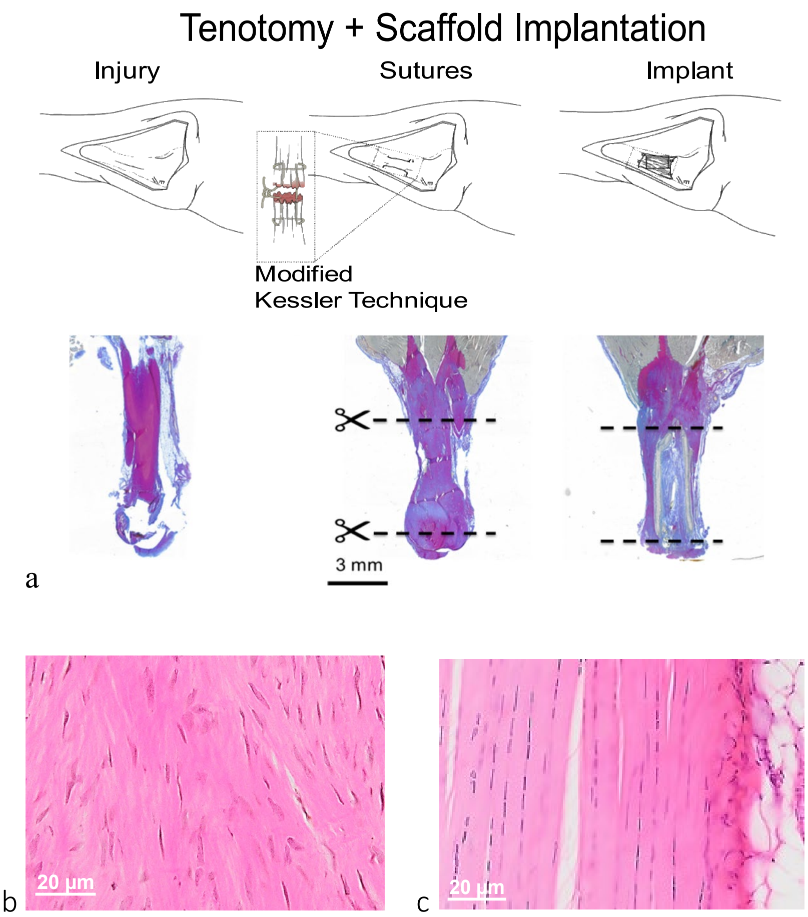


**Supplementary Fig. | 9.** **a**, Schematic representation and representative Herovici's stain histological images of both the intact tendon and injured tendon, repaired with either suture or with an electrospun scaffold. **b**, H&E images of injured tendon sample (1 week) and **c**, intact tendon sample showing differences in cell morphology. Pink, tissue matrix; purple, cytoplasm; blue, nuclei (Scale bar 20 and 10 µm respectively).

After the injury, the tendon stumps were observed to retract, generating a gap that became filled with granulation tissue. The fibrous regenerating tissue was characterised by loose collagen fibre organisation and an overall heterogeneous texture after two, four and eight weeks, with increased cellularity and ingrowth of vessels, nerves, fat deposition and calcification (Fig. SI10). At 1-week post-injury, the newly formed tissue was observed to be composed of disorganised granulation tissue comprised of collagen III. The digital analysis shows that the distribution of the orientation of fibres was broad and disorganised. A circular colour map shows the correlation between orientation and colour code. The polarised image demonstrates that the deposited collagen is highly disorganised and shows poor birefringence, characteristic of collagen type III.

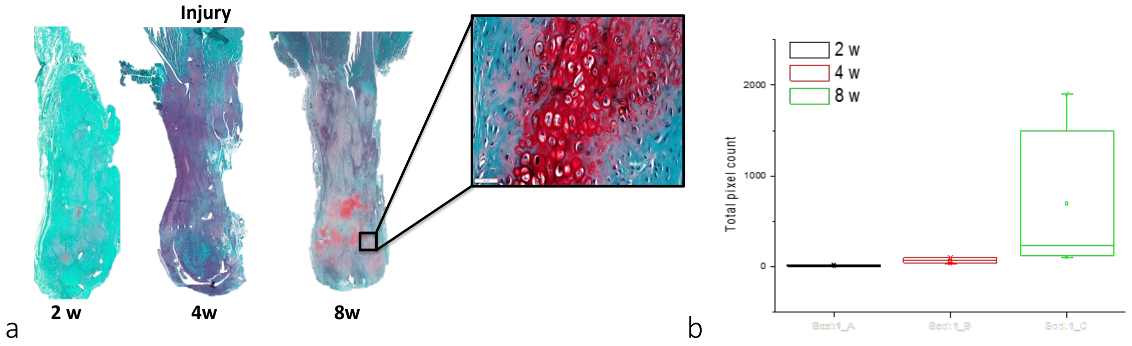


**Supplementary Fig. | 10. a**, Control (injury only) Safranin-O stained histological sections at two, four- and eight-weeks post-injury. After eight weeks, calcified areas in the mid-region of the tissue were observable. **b**, Quantification of the chondrification area for all samples after two, four- and eight-weeks post-injury. Histomorphometric analysis indicates a significant increase in chondrification eight weeks post-injury.

At two weeks post-injury, the tendon tissue composition was highly heterogeneous, possessing an increased cellular content and noted ingrowth of nerves and blood vessels relative to earlier time-points (Fig. SI11). At two weeks post-injury irregular voids were present between the large collagen type I fibres which were filled with granulation tissue and cells demonstrating a chondrogenic phenotype. At four weeks post-injury, a condensation of these cells showing a chondrogenic phenotype in both injury and piezo and non-piezo scaffold treated animals was observed. four weeks. This process, initiated by mechanical compression and the hypoxic conditions experienced by resident and infiltrating cells, was pronounced at the bone and tendon insertion areas. Indeed, by eight weeks post-injury endochondrial ossification was prominent in all static (cage) groups. Fig. SI11 shows the distribution of chondrocyte cells and cartilage nodules at the distal and in the mid-regions of the tendon granulation tissue following injury. As newly formed chondrocytes progressed in their maturation, the cartilage nodules were replaced by mineralised bone.

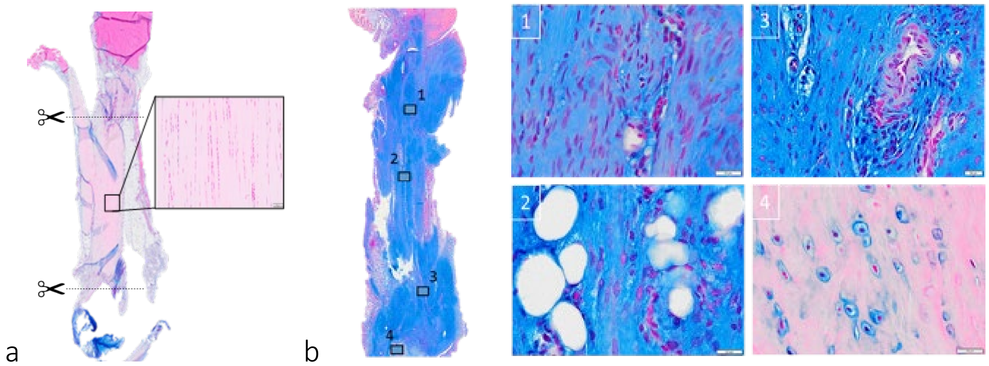


**Supplementary Fig. | 11**. Alcian blue stained histological sections of **a**, intact tendon showing aligned tenocytes and tendon ECM and **b**, 2 weeks post tendon transection repaired using a suture showing disruption of organised tissue and a significant presence of glycosaminoglycan (GAGs) in the matrix (blue). High-magnification images were subsequently from this section (indicated). 1) Upper region showing cells undergoing phenotype changes 2) Lipid deposition in the mid part of the injury 3) Vessel formation for cartilage formation at the lower region 4) Chondrocyte cells surrounded by their lacunae producing bone-specific matrix.

Following the inflammation phase (two weeks post-injury), tendon structure reorganisation was initiated, and areas of high cell density were observed close to the tendon stumps endings. Cartilage formation first appeared at the distal stump region. By four weeks post-injury, many cartilage nodules (Fig. SI11) were formed in the region of the granulation tissue close to the distal stump ending. Within this region, the chondrocytes matured into hypertrophic chondrocytes and mineralisation of the tissue was initiated. Several small cartilage nodules were present from four weeks both at the tendon stumps and throughout the granulation tissue. Progressively, cartilage nodules served as template for new bone formation via endochondral ossification (Fig. SI11), and ectopic bone formation resulted from hypertrophy of chondrocytes and blood vessel infiltration (Fig. SI11). To confirm ectopic bone tissue at eight weeks post-injury, micro CT scanning was employed in animals treated with suture, non-piezo and piezo scaffolds (cage). As shown in Figure SI12, mineralised tissue at the peri-implant region was observed, with the highest volume of calcified tissue observed in tissues repaired using the piezoelectric scaffolds under static conditions (cage).

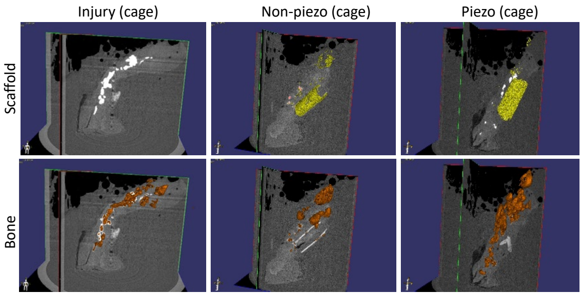

a

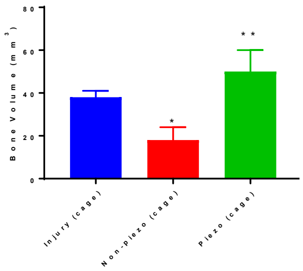

b

**Supplementary Fig. | 12. a**, Representative 3D micro CT reconstructions of injury in tendons treated with suture, non-piezoelectric and piezoelectric scaffolds eight weeks post tendon transection (cage). Mineralised tissue was observed in the distal part of the tendon corresponding to the bone insertion region. However, in animals treated the piezoelectric scaffolds, a significant amount of mineralised tissue was detected at the peri-implant region. **b**, Micro CT reveals quantitative differences in ectopic bone volume in tendon injuries treated with piezo and non-piezo scaffolds Analysis performed using one-way ANOVA with post-hoc Tukey HSD (N=3,*$p<0.05$ Non-piezo vs injury, ** $p<0.01$ piezo vs non-piezo).

In order to evaluate the degree of tendon repair and the effect of treadmill running on tendon ECM morphology, organisation and biochemistry, histomorphometric measurements on stained histological sections from three animals (5 slides in total per animal) were employed at eight weeks post-injury. A point-based composite scoring system was employed to grade tissue repair using parameters based on 1) the cell number relative to intact tendon 2) the tissue calcification relative to intact tendon 3) the tissue vascularisation and innervation relative to intact tendon 4) the extent of fat deposition relative to intact tendon 4) the fibre orientation relative to intact tendon 5) the cellular morphology relative to that of cells from to intact tendon.

**Supplementary Table 5**. A composite scoring system was employed for the histological evaluation of repair at eight weeks post-injury. Values are expressed in Mean ± SEM of scored points. 0-25 % (4 points); 26-50 % (3 points); 51-75% (2 points) and 76-100% (1 point).

| | Parameters | | | | | | |
|---|---|---|---|---|---|---|---|
| Group | Cell number | Calcified area | Vessels and nerves | Fat deposits | Fibre orientation | Cell shape (aspect ratio) | Total |
| Control | 4 ± 0 | 4 ± 0 | 3.5 ± 0.25 | 3.5 ± 0.5 | 3.75 ± 0.25 | 4 ± 0 | **23** |
| Injury | 1 ± 0.25 | 2 ± 0.5 | 3.25 ± 0.5 | 2.5 ± 0.5 | 1.5 ± 0.5 | 2.5 ± 0.5 | **12.75** |
| Injury (TR) | 1 ± 0.5 | 2.5 ± 0.25 | 2.5 ± 0.5 | 2.5 ± 0.5 | 2.5 ± 0.5 | 2.5 ± 0.5 | **13.5** |
| NonPiezo | 1 ± 0.75 | 2.25 ± 0.5 | 2.5 ± 0.5 | 2.5 ± 0.5 | 3.0 ± 0.5 | 2.5 ± 0.5 | **13.5** |
| NonPiezo (TR) | 2 ± 0.5 | 2.5 ± 0.5 | 3.25 ± 0.5 | 2.5 ± 0.5 | 3.25 ± 0.75 | 3 ± 0.5 | **16.5** |
| Piezo | 1 ± 0.75 | 1 ± 0.5 | 2.5 ± 0.5 | 2.25 ± 0.5 | 2.0 ± 0.75 | 2.5 ± 0.5 | **12.5** |
| Piezo (TR) | 1.5 ± 1 | 2.25 ± 0.5 | 2.5 ± 0.25 | 2.5 ± 0.5 | 3.25 ± 1.25 | 2.5 ± 0.75 | **15.0** |

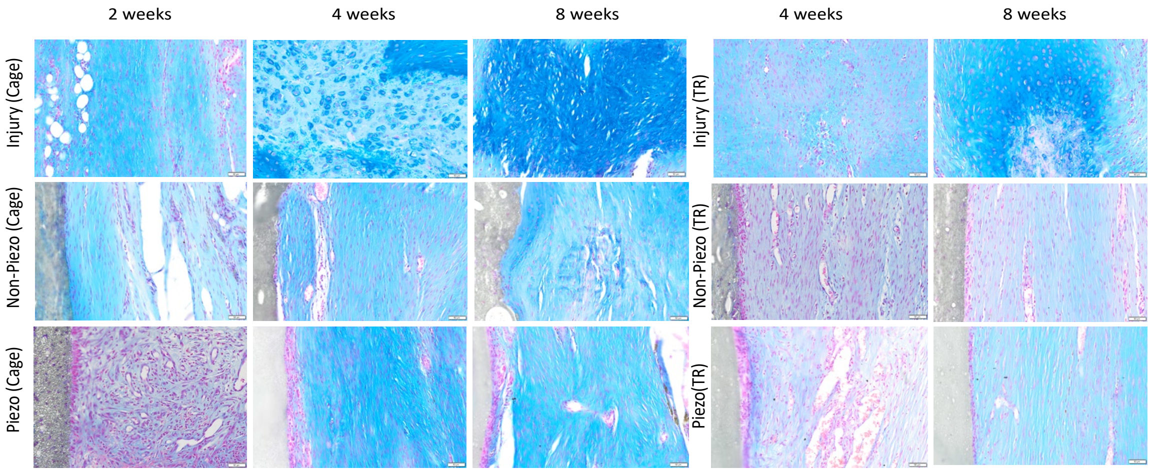


**Supplementary Fig. | 13**. Alcian-blue stained representative histological images indicating the role of electromechanical stimulation on proteoglycan deposition in animals subjected to static or treadmill running conditions. Tendon repaired with piezoelectric and non-piezoelectric scaffolds showed less intense proteoglycan (GAG) content. Conversely, transection injuries treated with a suture possessed highly disorganised tissues with increased cell proliferation, increased vasculature and fat deposition, and/or an increased GAG content. Scale bar is 200 µm.

During wound healing, early granulation tissue is produced by fibroblasts to act as a temporary scaffold for cell attachment and proliferation. Generally, granulation tissue is mainly comprised of collagen III, and low levels of collagen I. As the healing continues, the ratio of collagen I / collagen III becomes higher but generally never reaches its original values (around 1.4). As shown in Fig. 5b, a significant increase in the ratio of collagen type III to I was observed at four weeks for all TR groups yet returned to baseline eight weeks post-injury. Collagens are the main bulk constituent of tendon tissue. Collagen type I expression is associated with tissues under tensile loads and is the main protein responsible for the mechanical strength and durability of tendon tissue. An increase in the synthesis of collagen I over collagen III is a good indicator of tendon tissue maturation. During embryonic tendon tissue development, immature disorganised collagen III fibres mature into thick collagen, I fibres with strong birefringence and a highly aligned organisation. Conversely, Collagen II represents almost 80% of the total collagen content in cartilage and is specialised in supporting compressive forces. In tendon, collagen II appears in small amounts in the regions near the bone insertion. Conversely, fibrils of collagen III are always associated with collagen I.

In our study, we found that increased expression of the Piezo2 and TRPA1 (Fig. 5) ion channels at weeks two and four were observed in tendon tissues exposed to mechanical stimulation. The increased expression of these ion channels was positively correlated with the activation of the tenogenic pathways ERK/MAPK and FAK ($p<0.01$) at week four.

It is largely recognised MAPK/ERK signalling pathway is activated during exercise and is the link between exercise and adaptive changes in tendon composition. In addition, MAPK/ERK signalling pathway plays a central role in mediating cell division, migration and survival. The activation of the MAPK/ERK pathway was always observed in all (TR) groups relative to cage groups except for piezo (TR) at week eight ($p<0.01$) that showed significant dysregulation.

The upregulation of FAK signalling was significant only for non-piezo (TR) at week four and downregulated at week eight ($p<0.01$) (Fig. 5). The activation of the Smad-dependent pathway (TGF-β/BMP) was observed in (TR) groups at weeks four and eight compared to their corresponding static group ($p<0.05$) as shown in Fig. 5. The activation of BMP signalling pathways are involved in the homeostasis of the native tendon and during normal (healthy tendon) structural adaptation, collagen I to collagen III ratio was 1.4 and the main upregulated proteins were integrins β3 and β5, BMPR1A, Collagen I and V, Thrombospondin 4, Decorin and Tenomodulin.

The molecular pathways associated with the ectopic formation of bone after injury were studied, and significant upregulation in Wnt/β-catenin signalling was observed in caged animals treated with both piezoelectric and non-piezoelectric scaffolds at week four ($p<0.01$). In addition, it was observed that piezoelectrical scaffolds under static and dynamic (TR) conditions induced a transient activation of Wnt/β-catenin at week two ($p<0.01$) and eight ($p<0.01$) compared to the non-piezo (static) group that showed prolonged Wnt/β-catenin activation at weeks two, four and eight ($p<0.01$) (, SI14).

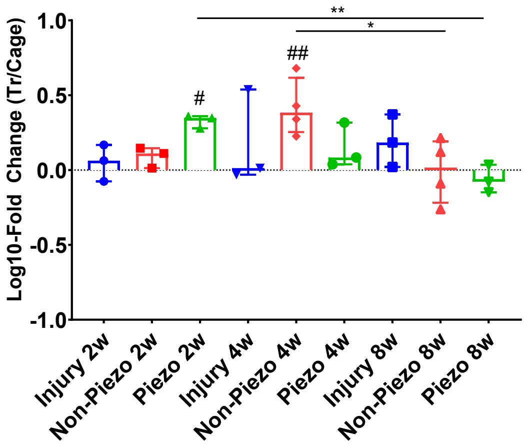


a)

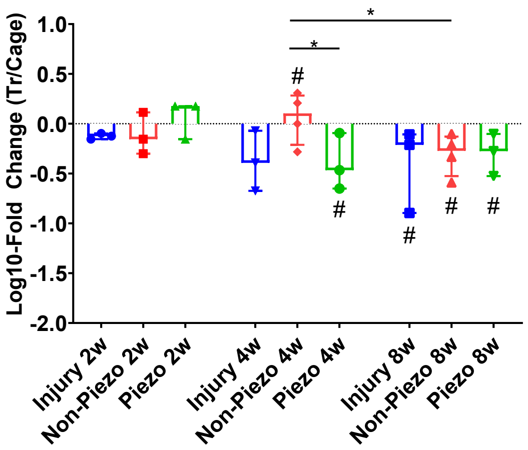


b)

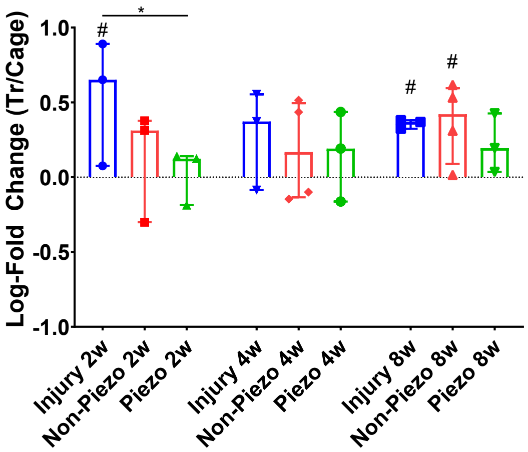


c)

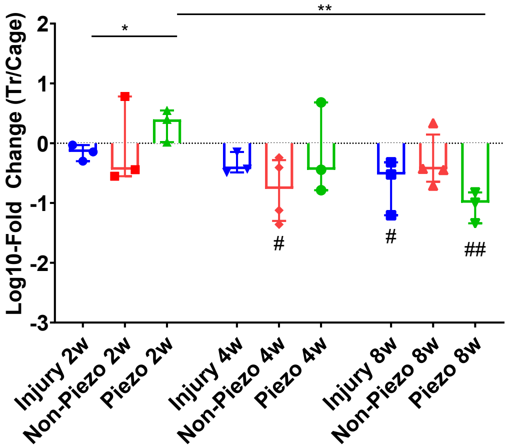


d)

**Supplementary Fig. | 14**. **a**, Protein Microarray data showing Log10-fold change in MAPK and **b,** FAK expression in tissues derived from TR animals relative to tissues derived from cage animals for each experimental group at weeks four and eight. Significance is shown between samples and week two (cage only). **c**, Protein Microarray data showing log-10 fold changes in BMP and **d,** Wnt expression in tissues derived from TR animals relative to tissues derived from cage animals for each group at weeks four and eight. Significance is shown between samples and week two (cage only). Analysis performed using Kuskal-Wallis (N=4) with post-hoc Dunn (*,# p<0.05. **, ## p<0.01).
